# Intrinsic mechanical sensitivity of auditory neurons as a contributor to sound-driven neural activity

**DOI:** 10.1101/2021.05.18.444619

**Authors:** Maria C. Perez Flores, Eric Verschooten, Jeong Han Lee, Hyo Jeong Kim, Philip X. Joris, Ebenezer N. Yamoah

**Affiliations:** Department of Physiology and Cell Biology, University of Nevada, Reno School of Medicine, Reno, NV 89557; Laboratory of Auditory Neurophysiology, Medical School, Campus Gasthuisberg, University of Leuven, B-3000 Leuven, Belgium.

## Abstract

Mechanosensation – by which mechanical stimuli are converted into a neuronal signal – is the basis for the sensory systems of hearing, balance, and touch. Mechanosensation is unmatched in speed and its diverse range of sensitivities, reaching its highest temporal limits with the sense of hearing; however, hair cells (HCs) and the auditory nerve (AN) serve as obligatory bottlenecks for sounds to engage the brain. Like other sensory neurons, auditory neurons use the canonical pathway for neurotransmission and millisecond-duration action potentials (APs). How the auditory system utilizes the relatively slow transmission mechanisms to achieve ultrafast speed and high audio-frequency hearing remains an enigma. Here, we address this paradox and report that the AN is mechanically sensitive, and minute mechanical displacement profoundly affects its response properties. Sound-mimicking sinusoidal mechanical and electrical current stimuli affect phase-locked responses. In a phase-dependent manner, the two stimuli can also evoke suppressive responses. We propose that mechanical sensitivity interacts with synaptic responses to shape responses in the AN, including frequency tuning and temporal phase-locking. The combination of neurotransmission and mechanical sensation to control spike patterns gives the AN a secondary receptor role, an emerging theme in primary neuronal functions.

## Introduction

The senses dependent on mechanosensation – hearing, balance, and touch – excel in speed and wide-ranging sensitivity among the sensory systems. The tactile sense has evolved beyond detecting simple mechanical stimuli to encode complex mechanical texture and vibration via unmyelinated and myelinated nerve endings and specialized mechanoreceptors(Delmas et al., 2011). For example, Pacinian corpuscles in mammalian skin encode vibrations up to ∼1000 Hz– more than an order of magnitude above the visual system’s flicker-fusion threshold, which sets the limit for stable viewing of computer screens and fluid motion in moving images. Transduction’s temporal acuity is directly translated into a neural code in such tactile receptors because transduction occurs in the same neural element that conducts the signal to the brain. This is different in the auditory and vestibular systems, where mechanosensation and messaging to the brain are subserved by separate cells. Their synaptic interface is a potential limit on temporal acuity.

Nevertheless, the vestibular system produces one of the fastest human reflexes, with a delay of only ∼5 millisecond(Huterer and Cullen, 2002). It is thought that non-quantal neurotransmission in the huge calyceal synapse between vestibular hair cells (HCs) and first-order neurons is an essential contributor to this speed(Eatock, 2018). Paradoxically, a specialized mechanism as observed in the tactile and vestibular systems, has not been identified in hearing, which is nevertheless the mechanosensitive system for which temporal acuity reaches its highest limits. For example, auditory brainstem neurons can reliably code tiny interaural time and intensity differences toward spatial hearing(Yin et al., 2019) over an enormous range of frequencies and intensities. The acknowledged mechanism for activation of classical action potentials (APs) in the primary auditory nerve (AN), also called spiral ganglion neurons (SGNs), is neurotransmission at the HC-ribbon synapse. Stereociliary bundles of auditory HCs convert sound-induced displacement and depolarization to neurotransmitter release unto AN afferents(Fettiplace, 2017; Fuchs et al., 2003; Roberts et al., 1988) with notable speed and temporal precision. The ribbon-type synapse is equipped to sustain developmentally regulated spontaneous activity (<100 Hz)(Levic et al., 2007), sound-evoked APs (<∼300 Hz), and phase-locked responses at auditory frequencies up to ∼5 kHz(Johnson, 1980). Phase-locking to sound stimuli is a nimble feature of the AN essential for sound detection, localization, and arguably for pitch perception and speech intelligibility(Peterson and Heil, 2020; Yin et al., 2019). How these response features remain sustained, despite the limits of presynaptic mechanisms of transmitter release to ATP-generation, synaptic fatigue, and vesicle replenishment(MacLeod and Horiuchi, 2011; Stevens and Wesseling, 1999; Yamamoto and Kurokawa, 1970), is not fully understood.

Responses of SGNs consist of multiple components, but the underlying mechanisms remain unresolved. Inner HC (IHC) depolarization results in increased excitatory postsynaptic current (EPSC) frequency. However, the synaptic current amplitude remains unchanged(Glowatzki and Fuchs, 2002; Grant et al., 2010). In response to low-frequency tones, the HC ribbon synapse triggers APs over a limited phase range of each sound cycle. Low- and high-intensity tones elicit phase-locked AN responses, which generate unimodal cycle histograms (i.e., response as a function of stimulus phase)(Johnson, 1980; Rose et al., 1967). In contrast, for some intermediate intensities, AN fibers fire APs at two or more stimulus phases, a phenomenon referred to as “peak-splitting” (Johnson, 1980; Kiang and Moxon, 1972). Moreover, a typical AN frequency tuning curve consists of two components(Liberman and Kiang, 1984) – a sharply tuned tip near the characteristic frequency (CF) and a low-frequency tail, which are differentially sensitive to cochlear trauma. Both of these observations suggest that more than one process may drive AN responses. Whereas the contribution of fast synaptic vesicle replenishment at the HC ribbon synapse is clear(Griesinger et al., 2005), the sustained nature of the IHC-AN synapse to sound(Glowatzki and Fuchs, 2002; Griesinger et al., 2005; Li et al., 2014) also raises the possibility that multiple processes sculpt AN responses.

Out of serendipity, we found that SGNs respond to minute mechanical displacement. Motivated by this finding, we hypothesized that SGN afferents actively sense organ of Corti movement and that this sensitivity, together with neurotransmission, shapes AN properties. The geometry of the course of the unmyelinated terminal segment of SGN dendrites towards the Organ of Corti (OC) suggests that this segment undergoes some degree of mechanical deformation in response to sound. An attractive feature of the SGN mechanical-sensitivity hypothesis is that it provides a straightforward substrate for one of two interacting pathways thought to generate peak-splitting and/or multi-component frequency tuning(Liberman and Kiang, 1984). Using *in vitro* simultaneous whole-cell recordings and mechanical stimulation, we show that adult SGNs are mechanically sensitive. Mechanical stimulation of the cell body (soma) elicits an inward current, which reverses at ∼0 mV. Mechanically-activated (MA) inward currents (I_MA_) and membrane voltage responses are sensitive to GsMTx4 peptide, a mechanosensitive channel blocker(Bae et al., 2011). Mechanical stimulation of the soma and dendrites elicits adapting, bursting, or non-adapting firing responses. Simultaneous sinusoidal stimulation with current injection and mechanical displacement alters firing rate and temporal coding. *In vivo*, single-unit recordings from AN fibers demonstrate that 2,3-dihydroxy-6-nitro-7-sulfamoyl-benzo[f]quinoxaline (NBQX) and Ca^2+^ channel blockers, potent inhibitors of synaptic transmission, suppress spontaneous APs, but sound-evoked APs persist at high intensities. These findings suggest that AN fibers’ intrinsic mechanical sensitivity contributes to sound-evoked activity and ill-understood features such as multi-component AN tuning curves and peak-splitting. Our findings blur the canonical distinction of the roles of sensory receptors and primary neurons and support the emerging notion that primary neurons may serve sensory receptor functions(Hattar et al., 2002; Woo et al., 2014).

## Results

### Mouse auditory neurons respond to stepped mechanical stimulation

**Figure 1a** shows the responses of adult mouse spiral ganglion neurons (SGNs) subjected to two different bath solution flow rates (0.5 and 3 ml/min). The neural activity at the highest flow rate is much larger and reflects the monotonously increasing relationship that saturates at 8 ml/min (Supplement 1, **S1**). In a small number of neurons (6 out of 110), increasing the flow rate produced a suppressive response (**S1**). Since SGN terminals are unmyelinated(Kim and Rutherford, 2016; Liberman, 1982), and subject to OC movement (OCM)(Chen et al., 2011; Jawadi et al., 2016; Karavitaki and Mountain, 2007), the finding raises the possibility of a direct mechanical pathway affecting AN responses in addition to synaptic transmission. To stimulate the unmyelinated dendritic terminals *in vitro*, SGNs were cultured on a polydimethylsiloxane (PDMS) substrate(Cheng et al., 2010). We displaced a single dendrite by substrate-indentation on this platform, using a fire-polished glass pipette driven by a piezoelectric actuator (**Sd, inset** **Fig. 1b**). Dendrite-substrate displacement sufficed to evoke membrane depolarization and APs at the recording patch-electrode (**R, inset** **Fig. 1b**). Soma-substrate or direct soma displacement was used for extended recordings and in voltage(V)-clamp experiments. In both stimulation configurations, rectangular- or ramp-displacements evoked either a sub-threshold membrane depolarization or APs, depending on the amplitude or ramp-velocity (slew rate) (**Fig. 1b, c**). When SGNs were interrogated with stepped mechanical displacement, 12 out of 32 neurons showed an increase in firing rate. In contrast, in some SGNs, including those with spontaneous activity (n = 10), an increase in mechanical displacement amplitude induced a suppressive response. In 10 out of 32 SGNs, interrogation with stepped mechanical displacement did not affect the AP firing rate (**S2b**).

**Figure 1.**
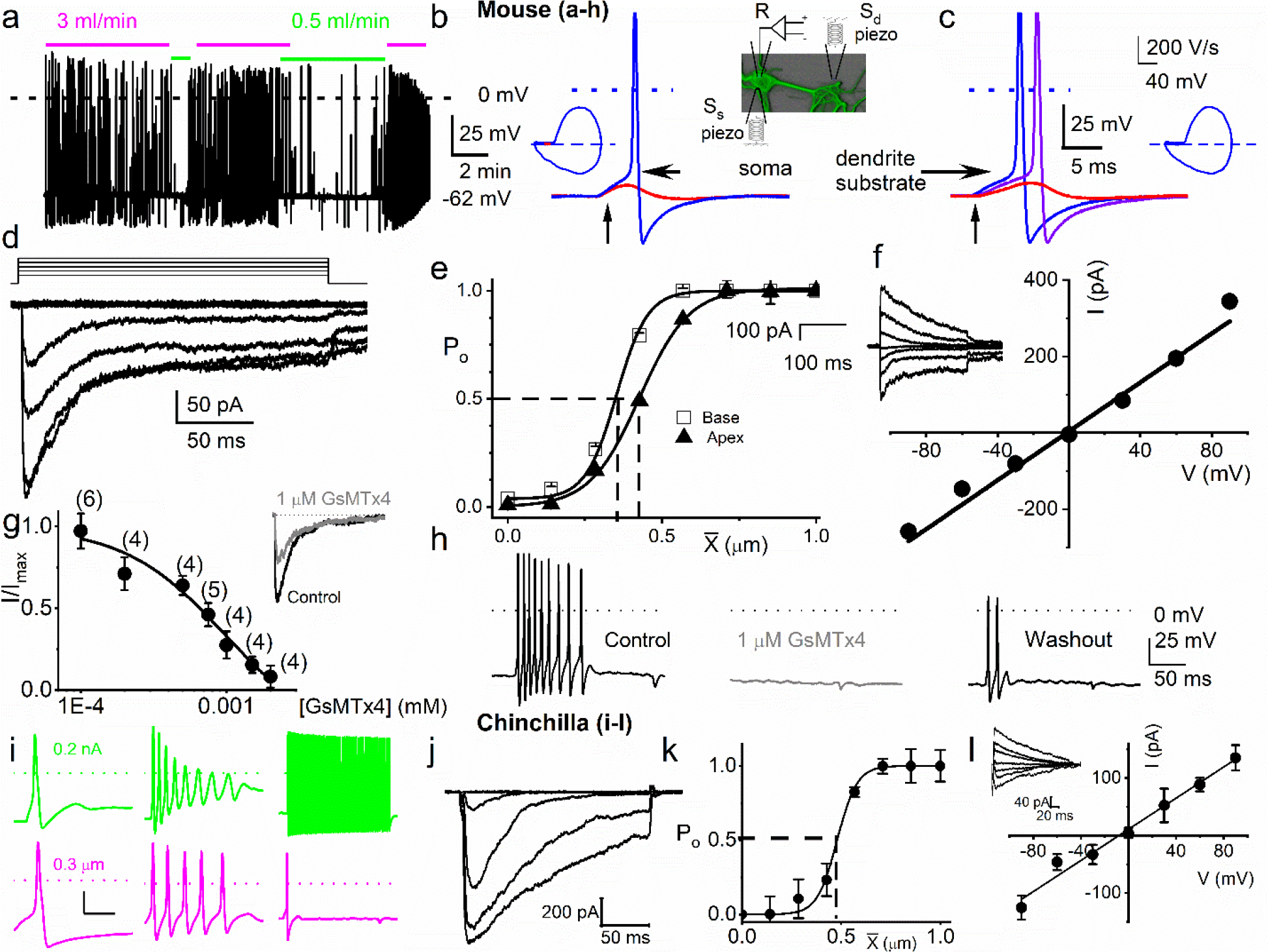
Mouse and chinchilla primary auditory neurons are mechanically sensitive. Mouse data (a-h) **a**, Responses of 8-week-old basal SGN to bath solution flow rate shown with colored bars (3 ml/min, magenta; 0.5 ml/min, green). V_rest_ = -62 mV. **b**, Mechanical displacement (20 ms rectangular-pulse injection) of apical SGN soma, sub-threshold (0.15 μm, red), and threshold (0.4 μm, blue) responses. Arrow indicates pulse initiation. Inset depicts the patch-clamp recording electrode (R) and stimulating probe (piezoelectric actuator represented as spring), placed ∼180° from the patch-electrode at the soma (Ss) or on the substrate to stimulate dendrites (S_d_). **c**, Responses of basal SGN to dendrite-substrate ramp-displacement at slew-rates (μm.ms^-1^) 0.1 (red), 0.5 (purple), and 1.0 (blue). Arrow indicates the time of pulse initiation. AP latency increased as the slew rate decreased (rate, 1-μm.ms^-1^, latency = 4 ± 0.1 ms; 0.5 μm.ms^-1^ = 6 ± 0.2 ms (*n* = 14). The corresponding phase plots (dV/dt versus V) of the responses are shown next to the traces. **d**, Current traces from displacement-clamp (X = 0 to 1.2 μm, step size ΔX = ∼0.24 μm) recordings at a holding voltage of -70 mV. Data are obtained from apical SGNs; for basal SGNs data, see **S2**. **e**, Summary data of displacement-response relationship of I_MA_ represented as channel open probability (P_o_) as a function of displacement, fitted by a single Boltzmann function. Data are from apical (**▴**) and basal (**□**) SGNs. The one-half-maximum displacements (X_0.5_) are indicated by the vertical dashed lines (X_0.5_ = 0.42 ± 0.01 μm (apical SGNs) and 0.35 ± 0.01 μm (basal SGNs), n = 14 for both). **f**. Average I-V relationship of MA currents. Inset shows traces of I_MA_ evoked at different holding voltages (-90 to 90 mV, ΔV = 30 mV, X = 0.4 μm). The regression line indicates a reversal potential of the MA current (E_MA_) of -1.4 ± 2.1 mV (cumulative data from apical SGNs, *n* = 17). The whole-cell conductance derived from the regression line is 3.2 ± 0.2 nS (*n* = 17). **g**, Dose-response relationship of I_MA_ as function of blocker GsMTx4, IC_50_ = 0.9 ± 0.2 μM (*n* = 4). Numbers of SGNs at different concentrations of GsMTx4 are shown in brackets. Inset shows I_MA_ in response to a 0.3 μm displacement at -70 mV holding voltage (gray trace is with 1 μM GsMTx4; black trace is control). **h**, APs from basal SGN evoked in response to 0.8 μm displacement of neurite-substrate (first black trace; V_rest_ = -59 mV; dashed line = 0 mV). Effect of 1 μM GsMTx4 (middle gray trace) and partial recovery after washout (last black trace; *n* = 5). **Chinchilla data (i-l): i**, Response properties (from left to right: fast, medium, and slow adapting) of three different chinchilla SGNs to 0.2 nA square-pulse injection (upper panel, green) and 0.3 μm soma displacement (lower panel, magenta). Note that the responses to mechanical displacement and current injection differed greatly. **j**, Current traces of an SGN soma (apical cochlear turn) in response to displacement-clamp stimuli from 0 to 1.2 μm in steps of ∼0.24 μm, at a holding potential of -70 mV. **k**, Displacement-response relationship of apical SGNs (n = 9), same as **e**, but for chinchilla; X_0.5_ is 0.49 ± 0.05 μm (vertical dashed line). **I**, Average I-V relationship of MA currents. Holding voltages range from -90 to 90 mV in increments of 30 mV; the inset shows I_MA_ traces at different holding voltages for a soma displacement of 0.4 μm. The regression line through the data points indicate an E_MA_ of -8.1 ± 5.5 mV (*n* = 5) and a whole-cell conductance of 1.4 ± 0.6 nS (*n* = 5).

From a holding potential (V_h_) of -70 mV, displacement evoked current (I_MA_) (**Fig. 1d**). The total I_MA_ amplitude ranged from 100-700 pA (426 ± 85 pA; *n* = 95). The I_MA_ shows a bi-exponential decay over time with a fast (τ_1_) and slow (τ_2_) time constant. For I_MA_ elicited with a 1.12 μm displacement, τ_1_ and τ_2_ were 3.6 ± 2.6 ms and 24 ± 5.7 ms (*n* = 17) for apical neurons, and 2.1 ± 1.1 ms and 17.0 ± 4.8 ms (*n* = 15) for basal neurons (**S3**), respectively. The displacement-response relationships of these apical and basal neurons (**Fig. 1e**), expressed as a channel open probability (P_o_), were fitted with a single Boltzmann function (black curves). The half-maximal activation displacements (X_0.5_) were 0.42 ± 0.01 μm (*n* = 14) and 0.35 ± 0.01 μm (*n* = 14) for I_MA_ from apical and basal neurons, respectively (**Fig. 1e**). Varying V_h_ and using a constant displacement (∼0.4 μm) yielded I_MA_ with a linear I-V relationship and a reversal potential (E_MA_) ∼0 mV (-1.4 ± 2.1 mV; *n* = 17), which is consistent with a non-selective cationic conductance (**Fig. 1f**). The I_MA_ in SGNs was sensitive to an externally applied MA channel blocker, GsMTx4(Bae et al., 2011). Application of 1 μM GsMTx4 decreased the current amplitude by ∼52% (52 ± 8%, *n* = 5; **Fig. 1g** inset). The half-maximal inhibitory concentration (IC_50_) obtained from the dose-response curve was 0.9 ± 0.2 μM (*n* = 4; **Fig. 1g**). Additionally, the application of 1 μM GsMTx4 completely abolished the dendrite displacement-evoked APs, which was partially reversible after washout (**Fig. 1h**).

### Response properties of chinchilla auditory and mouse vestibular neurons to stepped and sinusoidal mechanical stimulation

If the mechanosensory features of the SGNs are fundamental to auditory information coding, we would expect the phenomena to transcend species differences. Auditory neurons’ mechanical sensitivity was not restricted to the mouse: adult chinchilla SGNs were similarly, but not identically, responsive to mechanical stimulation, validating the potential physiological relevance across species. Shown are exemplary APs from chinchilla SGNs in response to the soma’s current injection and mechanical stimulation (**Fig. 1i**). The current elicited with mechanical displacement from -70-mV V_h_, and the summary of the corresponding displacement-response relationship was also fitted with a single Boltzmann function with X_0.5_ of 0.49 ± 0.05 (n = 9; **Fig. 1j** **& k**). The I-V relationship from chinchilla SGN I_MA_ produced E_MA_ = -8.1 ± 5.5 mV; n = 5 (**Fig. 1l**). The results suggest that I_MA_ in chinchilla SGNs share features in keeping with those observed in the mouse. A related issue is how specific the mechanical sensitivity of SGNs is relative to other mechanosensitive systems. Responses to mechanical steps of displacement were also observed in vestibular neurons (VN). The displacement-response relationship was approximated using a two-state Boltzmann function (**S4**). Compared to auditory neurons, mouse vestibular neurons (VN) were on average ∼3-fold in magnitude less responsive to stepped mechanical stimulation. Additionally, responses to high-frequency (> 10 Hz) mechanical stimulation were attenuated in VNs (**S4, see S6** for SGNs response to sinusoidal mechanical displacement). However, the mechanical responses of VNs were comparable to that reported for dorsal root ganglion (DRG) neurons(Finno et al., 2019; Viatchenko-Karpinski and Gu, 2016).

SGNs show a variety of responses to current injection. Current-evoked responses from 5-week old SGN range from fast to intermediate to slow adapting activity. Additionally, a fraction (5-10%) of SGNs are spontaneously active(Adamson et al., 2002; Wang et al., 2013) (**S5**). This diversity of responses is observed in adult apical and basal neurons(Lv et al., 2012). The question arises on how the response diversity to current injection relates to responsivity to mechanical stimulation. Apical and basal SGNs showed different response thresholds, with apical neurons showing higher sensitivity (**S5**). The response latency and excitability of SGNs were dependent on stimulus type (step vs. ramped pulse) and location (soma or dendritic substrate). For example, in fast-adapting SGNs, the first-spike latency increased from 4.0 ± 0.1 ms to 6.0 ± 0.2 ms (*n* = 14; *p* = 0.01) when the slew rate was reduced from 1 to 0.5 μm·ms^-1^; subsequent reduction in slew rate caused prolonged latency, subthreshold membrane depolarization, or failed responses (<0.001 μm·ms^1^; **S5**).

We used sinusoidal mechanical displacement as a proxy for *in vitro* sound stimulation at different frequencies, recording from mouse SGNs, and the responses were also variable (**S6**).

AP firing increased with increasing amplitude of mechanical stimulation (**S6**) and was phase-locked. In some SGNs (4 out of 11), the responses were attenuated with larger mechanical displacement. A train of varying frequencies of mechanical stimulation, in decade steps, shows the SGNs responded to and are phase-locked to specific frequencies up to 100 Hz, but responses were absent at 1000 Hz. In contrast, the response properties of SGNs reached 1000 Hz sinusoidal current injection. Responses to current injection were broader than mechanical stimulation (**S6)**.

### Amplitude and phase of combined current and mechanical sinusoidal stimulation affect response rate and timing

Because sound waves are converted into mechanical vibrations transmitted via the middle ear to the cochlea, which converts them into neural signals, the question arises whether the mechanical responses shown here play a role in hearing. As AN fibers traverse different compartments of the OC (osseous spiral lamina, basilar membrane) to innervate IHC, they lose their myelin sheath at the habenula perforata(Morrison et al., 1975) (**Fig. 2a**). The unmyelinated terminal is subject to sound-evoked displacement or pressure changes. Direct examination of potential synergistic or antagonistic effects between neurotransmission and mechanical signaling at or near the AN synaptic terminal is currently not technically possible; however, the interaction of their proxies of current injection and substrate vibration can be examined. We applied sinusoidal currents and mechanical displacements to produce more physiological stimuli that mimic sound-evoked neurotransmission and movement (see methods).

**Figure 2.**
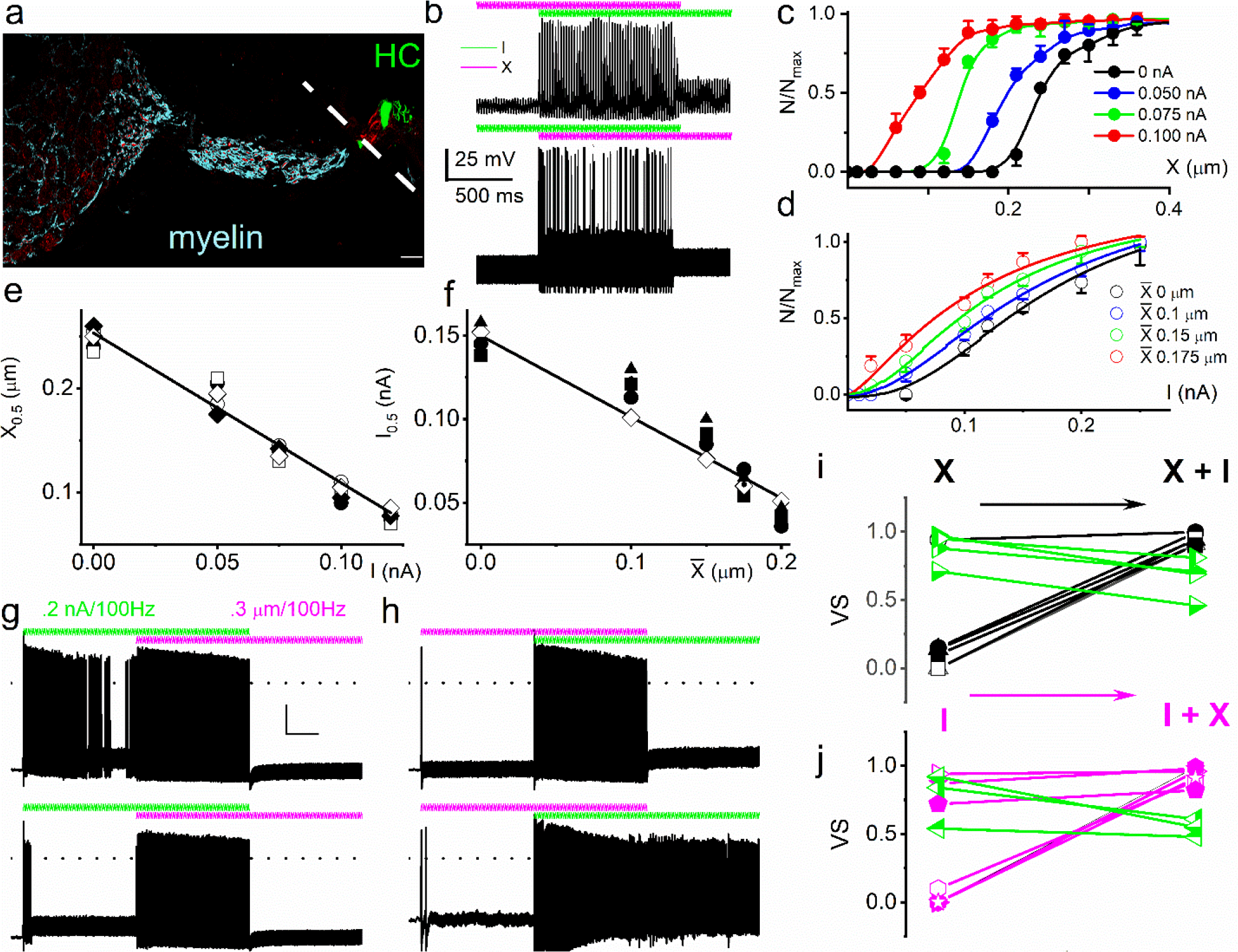
Mechanical stimulation of SGN cell-body increased sensitivity to current injection and affected phase locking. **a**, Horizontal section of whole-mount intact SGNs and HCs of a 5-week-old mouse cochlea, showing neuron (in red, anti-TUJ1) and myelin (in cyan, anti-myelin basic protein). The white dashed line shows the expected location of the basilar membrane. HCs are in green (anti-Myo7a). Note that the nerve terminals are devoid of myelin. Scale bar = 20 μm. **b**, (**Upper panel**) Voltage response to combined sub-threshold sinusoidal mechanical displacement (in magenta; 50 Hz, 0.1 μm, ∼2 s) and sub-threshold sinusoidal current injection (in green, 50 Hz, 0.05-nA, ∼2-s), delivered in-phase. Current injection and mechanical displacement overlapped for ∼1 s. Combined sub-threshold mechanical and current stimulation evoked APs. (**Lower panel**) We show conditions similar to the upper panel, but here sub-threshold current injection preceded sub-threshold mechanical displacement. Subthreshold stimuli, when paired, became suprathreshold. **c-d**, Spike-rate, normalized to maximum, as a function of displacement (**c**), and current injection (**d**). The leading stimulus was primed with a subthreshold sinusoidal current or displacement, respectively. **e-f**, The extent of sensitivity to current and displacement is represented as a plot of the half-maximum displacement (X_0.5_) and half-maximum current (I_0.5_) as a function of current and displacement. The slopes of the corresponding linear plots were, -1.4 μm/nA (n = 5) (**e**) and -0.5 nA/μm (*n* = 5) (**f**). **g**, (**Upper panel**) Slowly adapting SGN response to 0.2 nA injection at 100 Hz, overlapping with a pre-determined threshold (0.3 μm displacement at 100 Hz) mechanical stimulus. For this example, the vector strength (VS) transitioned from 0.82 to 0.98 upon combined (current plus mechanical displacement) stimulation (see Table 1 for summary data). (**Lower panel**) Moderately adapting SGN, stimulated with the same stimuli as the upper panel, transitioned from VS of 0.72 to 0.82 (Table 1). **h**, (**Upper and lower panels**) stimulation of fast adapting SGN in response to 100 Hz, a 0.3 μm displacement superimposed with 100 Hz 0.2 nA current injection. For the examples shown, the VS transitioned from unmeasurable to 0.94 for the upper panel and 0.9 for the lower panel. **i-j**, a summary of VS changes when either current or displacement is used as a primer for concurrent stimulation (Table 1 S6).

We find that the two stimulus modalities interact: mechanical and current stimuli, which are subthreshold by themselves, can be suprathreshold when combined (**Fig. 2b**). Since the exact relationship in amplitude and phase between synaptic and mechanical events in the cochlea is unknown, we explored different amplitude (**Fig. 2**) and phase (**Fig. 3**) relationships between current and mechanical stimulation. First, the interaction was determined using in-phase stimulation: SGNs were primed with sustained subthreshold current, and the displacement-response relationship was tested. The mechanical responsiveness increased with increased current injection, shifting the rate curves to lower displacement values (**Fig. 2c**). In parallel, increasing subthreshold mechanical displacement moved the rate curves to lower current values (**Fig. 2d**). The X_0.5_ derived from **Fig. 2c** as a current priming function has a linear relationship, with a slope of -1.4 μm/nA (**Fig. 2e**). The converse half-maximum current (I_0.5_; **Fig. 2d**), as a function of priming mechanical displacement, also shows a linear relationship, with a slope of -0.5 nA/μm (**Fig. 2f**). These orderly interactions between intrinsic mechanical responsiveness and electrical activity suggest that the increase in mechanical displacement with increasing sound intensity could have a monotonic effect on spike rate.

**Figure 3.**
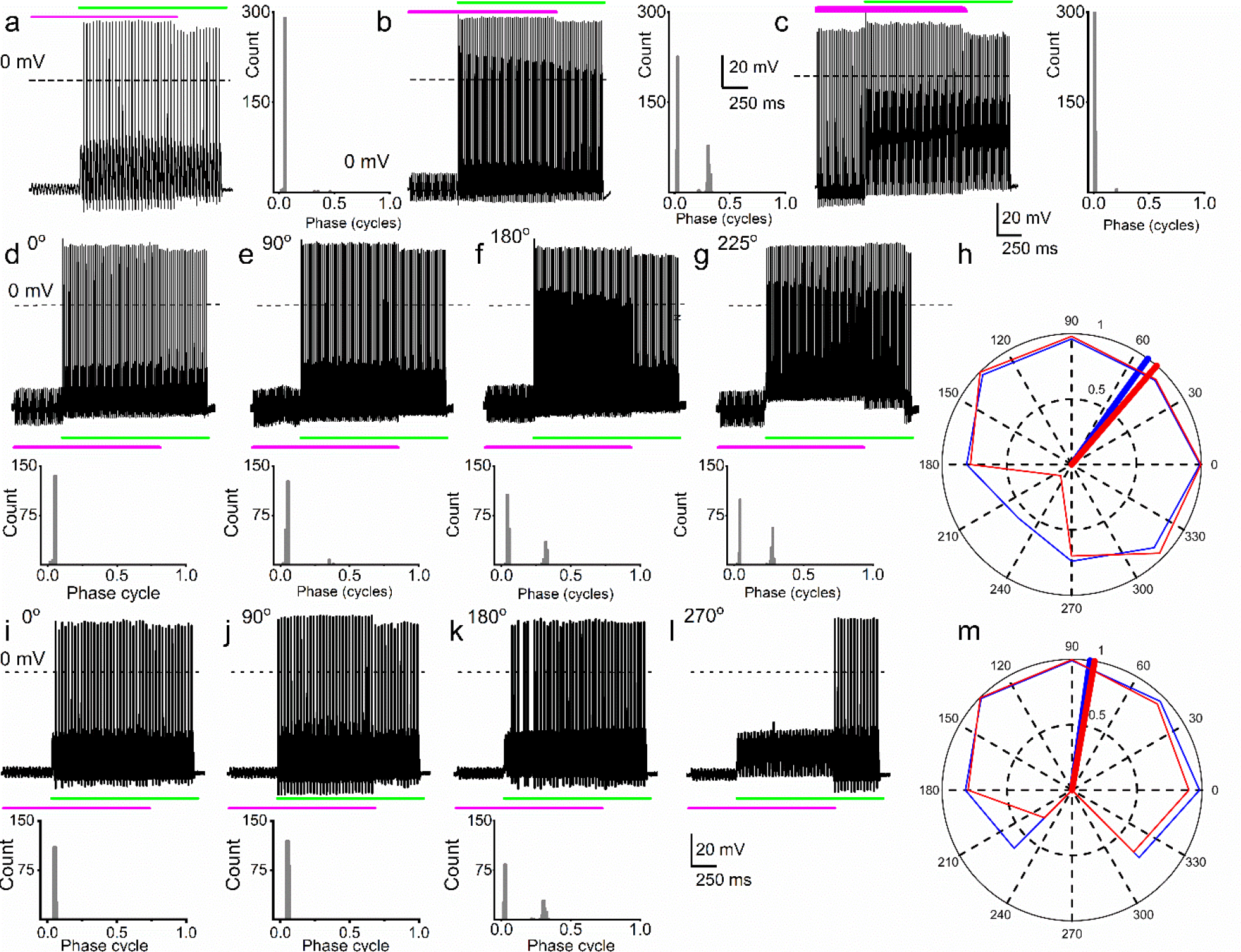
Simultaneous mechanical and electrical stimulation of SGNs can generate peak-splitting and rate suppression. **a,** SGN response properties and corresponding cycle histograms generated by simultaneous sinusoidal mechanical sub-threshold displacement (0.1-μm, 50 Hz; magenta) and sinusoidal current injection (0.2 nA, 50 Hz; green). The relative phase angle of the two stimuli was at 0°. The amplitude criterion of a valid AP was 0 mV (dashed line). **b, c.** The cycle histograms transitioned to two peaks and back to one peak as the amplitude of mechanical displacement increased to 0.2 μm (b) and 0.3 μm (c). **d-g**, Response properties and corresponding cycle histograms for the same SGN as in **a-c**, but using different relative phase angles (0°, 90°, 135°, 180°) between mechanical displacement (0.15 μm) and current injection (0.3 nA). At 180° and 225° phase angles, two peaks are evident in the cycle histograms shown in the panels below. h, Polar plot summarizing data averaged over 3 stimulus repetitions derived from the SGN illustrated in d-g. Different relative stimulus phase-angles are shown with the angular scale. The average rate is shown in blue and is normalized to its maximum; VS is shown in red. The solid blue and red line indicate the average angle for maximal firing and phase-locking: the close alignment of the two lines shows that a high firing rate is accompanied by strong phase-locking, and *vice versa* that peak-splitting is associated with low firing rates. **i-l**, Phase angle-dependent response reduction (180°) and suppression (270°) during combined stimulation. **m**, polar plot as in h but for the neuron illustrated in **i-l**.

The effect of combined stimulation on phase-locking was studied with in-phase sinusoidal current and mechanical displacement overlapped for ∼2 s. **Figure 2g** (upper panel) shows a slowly adapting SGN response to current stimulation (6-week-old basal SGN) with a firing rate of 36 ± 5 spikes/s, which increased to 50 ± 3 spikes/s upon paired mechanical stimulation applied to the soma. The vector strength (VS)(Goldberg and Brown, 1969) to the 100 Hz current injection alone was 0.82 ± 0.05 and increased with paired stimulation to 0.98 ± 0.01 (*p* = 0.01). For the moderately adapting SGN (**Fig. 2g** lower panel; 5-week-old apical SGN), the VS increased from 0.72 ± 0.06 to 0.82 ± 0.07 (*p =* 0.04) after paired stimulation. The converse paradigm, where mechanical stimulation preceded combined current and mechanical stimulation, is illustrated for two fast adapting SGNs. They showed a significantly increased AP firing rate when the two stimuli overlapped (**Fig. 2h**), with high VS values (0.94 and 0.9). For this set of experiments (28 SGNs) in which the pre-paired stimulation yielded non-zero VS (see Methods), 21 SGNs (75%) showed a significant increase in VS (*p* < 0.05). In the remaining 7 SGNs (25%), the VS was reduced (**Fig. 2i-j**, shown in green symbol and line). The summary data show that combined current and mechanical stimulation tended to alter the VS, suggesting the two stimuli interact to shape the response properties of SGNs (**Fig. 2i-j**, **Table 1**). When the response to mechanical stimulation was reduced by application of GsMTx4 (1 μM), dual current and displacement-responses resulted in a reduced VS (**S7, Table 1**).

**Table 1a.** Summary of changes in firing rate (R) and VS using in-phase current and mechanical stimulation. The significance level of VS is given in the column with *p* values obtained with the Rayleigh test for uniformity.(Mardia, 1972)

| SGN | Current (I)<br>R (spike/s)<br>VS | | Rayleigh<br>$p$ | Mech stim (X)<br>R (spike/s)<br>VS | | Rayleigh<br>$p$ | I + X<br>R (spike/s)<br>VS | | Rayleigh<br>$p$ |
| --- | --- | --- | --- | --- | --- | --- | --- | --- | --- |
| 13830002 | 0.5 | NA | NA | 0 | NA | NA | 40 | 0.96 | 0.0013 |
| 13830003 | 0 | NA | NA | 0 | NA | NA | 40 | 0.91 | 0.003 |
| 13903009 | 50 | 0.97 | 0.01 | 0 | NA | NA | 74 | 0.95 | 0.001 |
| 13903030 | 7.5 | 0.98 | 0.02 | 2.5 | 0.14 | 0.11 | 95 | 0.94 | 0.011 |
| 13903043 | 21 | 0.96 | 0.01 | 14 | 0.95 | 0.02 | 99 | 0.99 | 0.0003 |
| 13903044 | 11.5 | 0.95 | 0.01 | 51 | 0.98 | 0.01 | 100 | 0.99 | 0.0086 |
| 13903029 | 23 | 0.94 | 0.04 | 0 | NA | NA | 89 | 0.96 | 0.0002 |
| 13903010 | 49 | 0.98 | 0.01 | 0.5 | NA | NA | 69 | 0.94 | 0.0012 |
| 13903058 | 45 | 0.98 | 0.01 | 12 | 0.97 | 0.02 | 96 | 0.97 | 0.0003 |
| 13903059 | 24.5 | 0.94 | 0.03 | 50.5 | 0.98 | 0.01 | 92 | 0.96 | 0.002 |
| 13904007 | 0.5 | NA | NA | 0 | NA | NA | 99.5 | 0.99 | 0.009 |
| 13904008 | 0 | NA | NA | 22.5 | 0.94 | 0.03 | 100 | 0.997 | 0.0001 |
| 13904023 | 1.5 | 0.15 | 0.43 | 0 | NA | NA | 99.5 | 0.99 | 0.0002 |
| 13904024 | 0 | NA | NA | 1 | NA | NA | 100 | 0.96 | 0.003 |
| 13904037 | 2 | 0.1 | 0.67 | 0 | NA | NA | 55 | 0.89 | 0.008 |
| 13904038 | 0 | NA | NA | 2.5 | 0.15 | 0.32 | 50.5 | 0.995 | 0.008 |
| 13909008 | 0.5 | NA | NA | 0 | NA | NA | 64 | 0.96 | 0.012 |
| 13909009 | 0 | NA | NA | 1.5 | 0.12 | 0.52 | 62 | 0.96 | 0.013 |
| 13909024 | 36 | 0.87 | 0.02 | 0 | NA | NA | 50 | 0.98 | 0.011 |
| 13909025 | 0 | NA | NA | 0.5 | NA | NA | 50.5 | 0.95 | 0.012 |
| 13909040 | 0 | NA | NA | 0 | NA | NA | 16 | 0.94 | 0.007 |
| 13909041 | 0 | NA | NA | 0 | NA | NA | 13 | 0.75 | 0.052 |
| 13913008* | 45.5 | 0.72 | 0.04 | 0 | NA | NA | 43 | 0.82 | 0.036 |
| 13913009* | 35 | 0.87 | 0.03 | 1 | NA | NA | 50.5 | 0.9 | 0.027 |
| 14900012* | 18 | 0.76 | 0.02 | 3 | 0.15 | 0.34 | 64.5 | 0.95 | 0.011 |
| 14903018* | 0.5 | NA | NA | 5 | 0.2 | 0.36 | 31.5 | 0.87 | 0.021 |
| 14903021* | 1.5 | 0.1 | 0.25 | 0.5 | NA | NA | 45.5 | 0.91 | 0.010 |
| 14903083 | 32.5 | 0.68 | 0.02 | 0.5 | NA | NA | 44.5 | 0.83 | 0.008 |
| 15800101 | 0.5 | NA | NA | 3 | 0.1 | 0.31 | 72.5 | 0.96 | 0.006 |
| 15800110 | 56 | 0.95 | 0.01 | 1 | NA | NA | 62 | 0.93 | 0.071 |
| 15800099 | 32 | 0.94 | 0.02 | 0 | NA | NA | 70.5 | 0.96 | 0.412 |
| 15810007 | 56 | 0.98 | 0.01 | 3.5 | 0.2 | 0.14 | 58 | 0.97 | 0.207 |
| 15800291 | 0 | NA | NA | 5 | 0.25 | 0.08 | 94.5 | 0.99 | 0.001 |
| 15800292 | 0.5 | NA | NA | 3 | 0.10 | 0.56 | 56.5 | 0.92 | 0.005 |
| 15800293 | 3 | 0.15 | 0.09 | 1 | NA | NA | 95.5 | 0.99 | 0.0001 |
| 15800295 | 2.5 | 0.1 | 0.27 | 3 | 0.1 | 0.53 | 68.5 | 0.89 | 0.004 |
| 15800301 | 1 | NA | NA | 8 | 0.3 | 0.18 | 46.5 | 0.96 | 0.0002 |
| 15800308 | 48 | 0.87 | 0.02 | 1 | NA | NA | 92 | 0.98 | 0.003 |
| 15800319 | 1 | NA | NA | 7 | 0.25 | 0.11 | 89.5 | 0.94 | 0.003 |
| 15800320 | 0 | NA | NA | 5 | 0.3 | 0.11 | 49.5 | 0.91 | 0.003 |
| 15800401 | 16.5 | 0.71 | 0.02 | 17 | 0.55 | 0.03 | 56.5 | 0.89 | 0.021 |
| 15800402 | 7 | 0.25 | 0.05 | 1 | NA | NA | 42.5 | 0.91 | 0.025 |
| 15800511 | 3 | 0.1 | 0.18 | 1 | NA | NA | 40 | 0.94 | 0.008 |
| 15800513 | 5 | 0.2 | 0.09 | 6 | 0.14 | 0.17 | 52 | 0.94 | 0.006 |
| 15800515 | 1 | NA | NA | 94.5 | 0.98 | 0.003 | 89.5 | 0.99 | 0.002 |
| 16900101 | 0 | NA | NA | 97.5 | 0.95 | 0.01 | 99.5 | 0.998 | 0.002 |
| 16900107 | 5 | 0.2 | 0.05 | 3 | 0.1 | 0.21 | 75.5 | 0.96 | 0.001 |
| 16900111 | 9 | 0.3 | 0.05 | 5 | 0.15 | 0.11 | 98 | 0.995 | 0.007 |
| 16900123 | 1 | NA | NA | 6 | 0.12 | 0.24 | 72.5 | 0.96 | 0.0002 |
| 16900127 | 19 | 0.74 | 0.01 | 1 | NA | NA | 92 | 0.95 | 0.008 |
| 16900131 | 9 | 0.25 | 0.03 | 1 | NA | NA | 45.5 | 0.75 | 0.037 |
| 16900139 | 5 | 0.2 | 0.12 | 4 | 0.1 | 0.67 | 56 | 0.9 | 0.009 |

SGN = spiral ganglion neuron, R (spike rate) = spike/s, VS = vector strength, \* substrate dendrite displacement, NA = not applicable.

**Table 1b.** Effects of GsMTx4 on firing rate and VS

| SGN | Current (I) |  | Mech stim (X) |  | I + X |  |
| --- | --- | --- | --- | --- | --- | --- |
|  | R (spike/s) | VS | R (spike/s) | VS | R (spike/s) | VS |
| 19830002 | 3 | 0.15 | 1.5 | 0.1 | 41 | 0.90 |
| GsMTx4 | 3 | 0.15 | 0 | NA | 5 | 0.24 |
| 19830002 | 45.5 | 0.74 | 1 | NA | 48 | 0.86 |
| GsMTx4 | 49.5 | 0.75 | 0 | NA | 42 | 0.72 |
| 19830043 | 9 | 0.62 | 5 | 0.3 | 31 | 0.89 |
| GsMTx4 | 11 | 0.62 | 0 | NA | 13 | 0.58 |
| 19830059 | 7 | 0.38 | 14 | 0.64 | 45 | 0.89 |
| GsMTx4 | 5 | 0.37 | 0 | NA | 11 | 0.37 |

Not only are the amplitude and phase of the OCM relative to IHC stereociliary bundle displacement not known: they will also vary, depending on stimulus parameters(Cooper and Rhode, 1995). Therefore, the effects of sinusoidal displacement and current injection at different phase angles and amplitudes were tested. The top row of **Fig. 3a-c** shows SGN responses to combined mechanical (magenta) and current (green) stimulation – the 50 Hz stimuli are in-phase but are changed in relative amplitude by increasing the displacement amplitude in steps of 0.1 μm. Combining subthreshold (0.1 μm) mechanical stimulation with a suprathreshold (0.2 nA) current triggered a highly phase-locked response, as illustrated by the cycle histogram (**Fig. 3a**, right panel), which shows the occurrence of spiking as a function of stimulus phase. The increase of the mechanical stimulus amplitude to 0.2 μm causes spikes to appear at a different phase, giving rise to a second peak in the cycle histogram, reminiscent of the *in vivo* phenomenon of peak-splitting(Kiang and Moxon, 1972; Liberman and Kiang, 1984). With a further increase in amplitude to 0.3 μm, the cycle histogram is again unimodal (**Fig. 3c**). Peak-splitting with paired mechanical-current stimulation was also observed in chinchilla SGNs (**S9**). On rare occasions (2 out of 48 SGNs), three APs were evoked per cycle, as shown for a chinchilla SGN (**S8**). To test the effect of the relative phase of the mechanical stimulus, it was changed in 45° increments with respect to the current while measuring response rate and phase. Peak splitting could be observed at some phase angles (**Fig. 3f,g**) and not at others (**Fig. 3d, S9**). To obtain a visual overview of all phase angles at which peak-splitting occurred, we created polar graphs showing VS (red) and spike rate (blue) for each stimulus phase angle (**Fig. 3h,m, S10**). Because a drop in VS necessarily accompanies peak-splitting, its occurrence is reflected in reduced VS values (e.g., at 225° in **Fig. 3h**). In stark resemblance to observations *in vivo*, there was a decrease in spike rate at the phase angles where peak-splitting occurred, illustrated by the alignment in average vectors for rate and VS (**Fig. 3h,m, S10**: thick red and blue lines). *In vivo*, this phenomenon has been called “Nelson’s notch”(Kiang and Moxon, 1972; Liberman and Kiang, 1984), and until now had not been recapitulated *in vitro*. This decrease could result in spike rates lower than those elicited by the single-stimulus conditions – e.g., in **Fig. 3l**, the spike rate during overlap is lower than due to current injection only. Thus, paired stimuli can not only generate a monotonic increase in spike rate (**Fig. 2c-f**) but can also cause response suppression. Further examples of the correlation between spike rate and VS are shown in **S10** from 11 series of measurements with relative phase between current and mechanical stimuli as the independent variable.

### Auditory nerve responses in the absence of synaptic transmission

To examine whether AN fibers are influenced directly by mechanical displacement *in vivo*, we administered blockers of synaptic transmission through the cochlear round window (RW) while recording from single fibers. Reports have shown two response regions in frequency tuning curves: a “tip” of sharp tuning observed at low sound levels, and a “tail” of coarse tuning (usually below the CF) at high sound levels(Kiang et al., 1986). These two regions are differentially vulnerable to experimental manipulations(Kiang et al., 1986). The inherent mechanical sensitivity of AN fibers demonstrated *in vitro* is observed for substrate motion at sub-micrometer displacements (< 100 nm, e.g., **Fig. 2c**). But note that temporal effects can occur at subthreshold displacements, i.e., that do not trigger spikes themselves, e.g., **Fig. 3**). Cochlear displacements of this magnitude have been measured *in vivo*, particularly towards the cochlear apex and at high sound pressure levels (SPLs), where frequency tuning is poor(Cooper and Rhode, 1995; Lee et al., 2015; Lee et al., 2016a). The *in vitro* results show interactive effects of current injection and mechanical stimulation (**Figs. 2,3**), so it is difficult to predict the response that would remain when one modality (synaptic-evoked spiking) is abolished. Nevertheless, the expectation is that *in vivo*, in the presence of synaptic blockers, some degree of response will persist at high SPLs, both in normal and traumatized cochleae.

We first characterized frequency tuning in single AN fibers during one or more electrode penetrations through the AN, recorded at the internal auditory meatus. After a sample of frequency tuning curves was obtained (**Fig. 4a**), sufficient to record a frequency tuning profile as a function of recording depth, NBQX was applied through the round window. NBQX is a competitive antagonist of the ionotropic glutamate-receptor, which blocks HC-AN synaptic transmission(Grant et al., 2010). The AN recording was then resumed with the same micropipette, left *in situ* during drug injection. NBQX application progressively decreased spontaneous- (**Fig. 4d**) and sound-evoked (**Fig. 4b**) spiking activity. Because these recordings are inherently blind, AN fibers without spontaneous activity can be detected only using brief high-intensity sound “search” stimuli (see Methods). Figures **4a-b** show tuning curves recorded before and after NBQX administration. Notably, after NBQX application and at low recording depths, fibers were excitable only at high SPLs. Tuning in these fibers was broad and shallow, with the lowest thresholds at low frequencies (**Fig. 4c**). We surmised that these responses persisted as a result of the presence of intrinsic mechanical sensitivity. However, interpretation of the response is hampered by the difficulty of knowing to what frequency range the fibers were tuned before application of the blocker. Because the high-threshold, coarsely-tuned responses were observed at recording-depths where, pre-NBQX, neurons were tuned to high frequencies, it can be reasonably inferred that these responses originated from fibers innervating the cochlear base. Fibers recorded at depths > 200 µm were tuned to frequencies < 1 kHz, both pre-and post-drug applications (**Fig. 4c**). Limited intra-cochlear diffusion of the drug likely accounts for lesser drug effects at these CFs.

**Figure 4.**
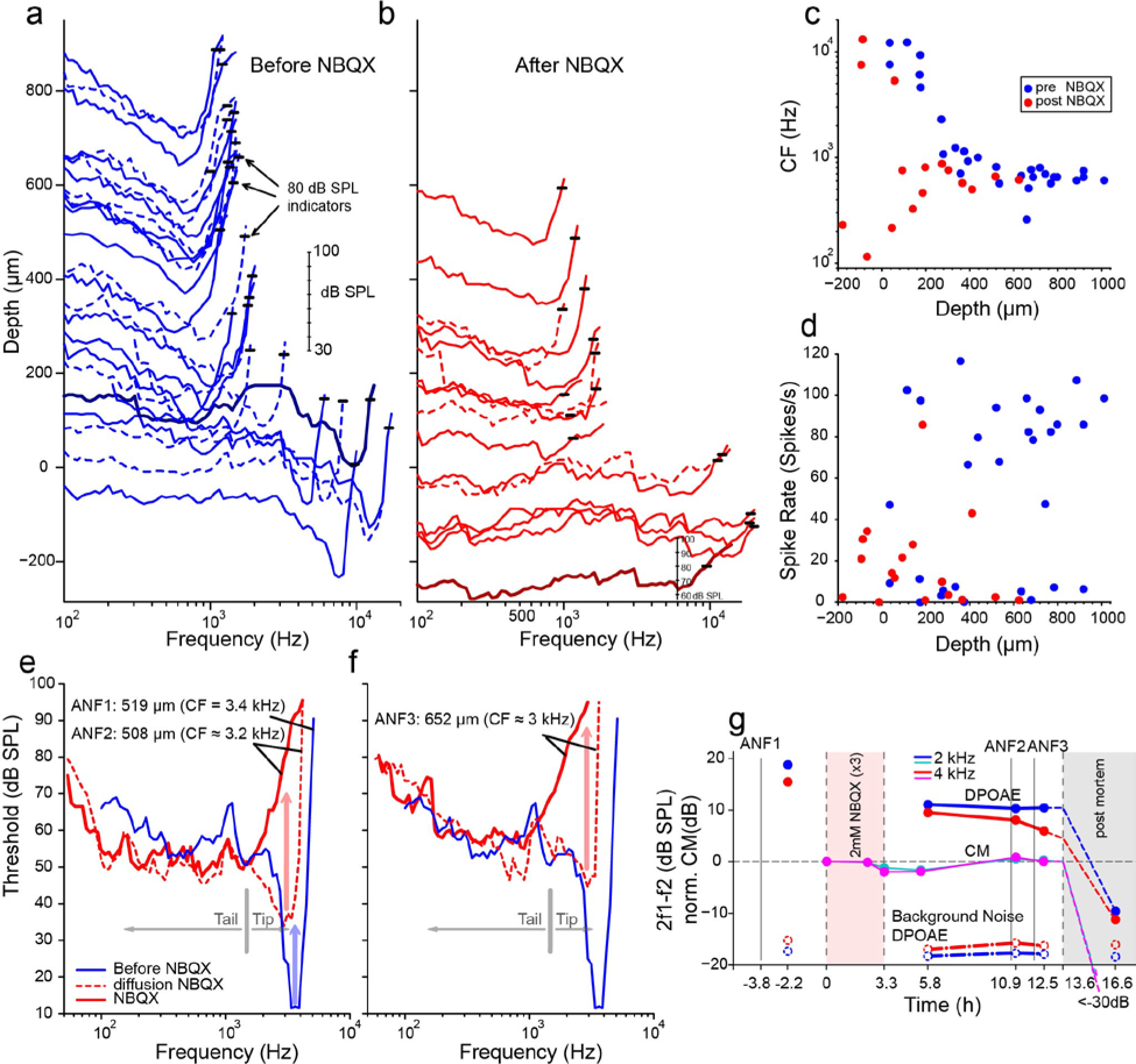
Effects of blocking synaptic transmission on AN activity *in vivo*. **a-b**, Frequency threshold tuning curves of AN fibers from a single chinchilla, obtained at different recording depths in the nerve, before (blue) and after (red) round window application of NBQX. Each curve shows the threshold (in dB SPL) of a single AN fiber over a range of frequencies: the curves are anchored to the y-axis (recording depth) by the small black horizontal bar at 80 dB SPL. The dB scale in **a** applies to all traces in **a,b**. Dashed lines are sometimes used to disambiguate traces. After applying NBQX, frequency tuning vanishes at the shallow recording depths (toward the bottom of **a** and **b**), where high-frequency fibers were found before applying NBQX. Fibers tuned to lower frequencies, found at greater depth (toward the top of **a** and **b**), are less affected, likely due to limited diffusion of NBQX to the apical turns. **c**, Characteristic frequencies as a function of recording depth (CFs, frequency of lowest threshold) before and after application of NBQX in the same cochlea. In the initial recording sessions, before the application of NBQX (blue symbols), high CFs dominate at shallow recording depths, while at depths > 200 µm CFs are < 1 kHz. After application of NBQX (red symbols), the lowest thresholds of superficial fibers are predominantly at low frequencies (< 1 kHz). **d**, spontaneous rates as a function of recording depth. Formatting as in **c**. **e**, the effect of NBQX on repeated measurement of a threshold tuning curve of AN fiber 2 (ANF2) recorded initially (red dashed) and after several minutes (red solid), compared to a tuning curve of a fiber (ANF1) with comparable CF recorded before NBQX application (blue). **f**, same as in **e**, but for a different AN fiber (ANF3). **g**, monitoring of the mechanical cochlear state during the experiment in **e** and **f** at two relevant frequencies, using Distortion Product OtoAcoustic Emissions (DPOAEs) and compound receptor potentials (cochlear microphonic, CM). Amplitudes of DPOAEs are shown in dB SPL; CM amplitudes are normalized to the level measured before application of NBQX (measurements **1-2**). CM and DPOAEs are relatively stable after the NBQX injection but vanish postmortem. The measurement noise floor (dashed lines) is stable over the entire experiment.

While the data in **Fig. 4a, b** are consistent with previous reports showing differences in the vulnerability of low- and high-threshold components of AN threshold curves, two fibers shown in **Fig. 4e,f** provide a more direct indication that high-threshold, bowl-shaped tuning originated from basally located AN fibers. In these two fibers, the “tip” but not the “tail” region of the tuning curve disappeared during repeated measurement, such that only a bowl-shaped high-threshold response to low-frequency tones remained. For reference, **Fig. 4e,f** shows the tuning curve of a fiber with CF of 3.4 kHz (blue, spontaneous rate of 8.7 spikes/s) recorded at 519 µm depth before NBQX application; the other curves (**Fig. 4e,f**, red) were obtained post-NBQX. The dashed red curve in **Fig. 4e** is the tuning curve of a fiber recorded at a similar depth (508 µm) after the NBQX application. It reveals a tip at 3.2 kHz (spontaneous rate of 15.3 spikes/s) with an elevated threshold and little separation between threshold at the tip and tail. Presumably, the tuning curve before drug application was similar in shape to that of the fiber recorded at this depth pre-NBQX (blue curve). A second, subsequent measurement of the same fiber (solid line, **Fig. 4e**) shows a curve consistent with the tail measured initially but with the tip attenuated. A similar progression was measured for a second fiber, measured at a greater depth (652 µm, initial spontaneous rate = 23.3 spikes/s), and it showed a similar pattern of a tip near 3 kHz, which subsequently disappeared with a broad bowl-shaped curve remaining in a follow-up measurement (spontaneous rate = 15.9 spike/s). Combined, these data (**Fig. 4a-f**) reveal a difference in vulnerability between the nerve fiber’s tip and tail, consistent with different modes of activation in the two response regions.

It is conceivable that the loss of sensitive tips as in **Figs. 4b,e,f** does not reflect a loss of synaptic drive but rather a decline in cochlear health and sensitivity. Even if such a functional cochlear decline is present, we still expect NBQX to block synaptic transmission and eliminate or reduce sound responses. The observed remaining response is thus consistent with a contribution by intrinsic mechanical sensitivity of nerve fibers, independent of the source of the loss of the tip of frequency tuning. Moreover, as assessed by cochlear emissions and cochlear microphonics monitored throughout the experiment, the response in **Figs. 4a-f** were from healthy cochleae. Acoustic distortion products elicited by tones spanning the CF range of the two fibers illustrated in **Fig. 4e,f**, remained high after injection of NBQX, as did the cochlear microphonic at similar frequencies (**Fig. 4g**). Both signals disappeared when the experiment was terminated by anesthetic overdose. These observations support the interpretation that, even though the OC’s underlying vibrations were intact, the tuning curve’s tip was abolished by the synaptic blocker because it only depends on synaptic input. Simultaneously, the tail is more resistant because it reflects a synaptic drive combined with intrinsic mechanical sensitivity.

It is also conceivable, albeit unlikely, that synaptic activation is qualitatively different at high- and low-sound levels so that a competitive blocker such as NBQX does not effectively block synaptic transmission at high-sound levels. The effects of nimodipine and isradipine L-type Ca^2+^ channel and synaptic exocytosis blockers(Huang and Moser, 2018; Rodriguez-Contreras and Yamoah, 2001) were also tested. **Figure S11a,b** shows tuning curves measured before and after the injection of nimodipine through the round window. There is a profound suppression of spontaneous activity for recording depths up to ∼500 µm, accompanied by very shallow tuning curves, often consisting of only tails (**S11b-d**). Deeper in the nerve, spontaneous rates and shapes of tuning curves are less profoundly affected, likely due to limitations in the diffusion of the drug to more apical cochlear regions. The application of isradipine produced similar effects.

## Discussion

AN fibers’ encoding responses are remarkably similar to other sensory neurons, with short-duration APs and inherently delayed neurotransmitter mechanisms(Glowatzki and Fuchs, 2002; Griesinger et al., 2005; Li et al., 2014). It has been a paradox how the AN utilizes conventional neural mechanisms to operate at 100- to 1000-fold faster time-scales with acute precision and reliability, compared to other neural systems(Hudspeth, 1997; Koppl, 1997). Puzzling auditory features such as multi-component frequency tuning and phase-locking may serve as gateways to gain mechanistic insights. Motivated by an unexpected finding that AN respond to minute mechanical displacement, an investigation was conducted to show whether, in addition to using established pathways for neurotransmission, type I SGNs are mechanically sensitive at their unmyelinated nerve endings.

It was discovered that simultaneous current injection (simulating the effect of neurotransmission) and mechanical displacement could interact to affect AN firing rates and phase-locked responses. Combined current injection and mechanical activation of the AN affects the phase angle at which spikes are fired and can generate peak-splitting, and both spike rate increase and suppression. *In vivo* recordings from chinchilla and cat demonstrate that blocking the AN activation pathway via neurotransmission does not suffice to eliminate sound-evoked AN spiking at high sound levels. Therefore, it is proposed that sound-evoked spiking in the AN, particularly at high-sound levels, is buoyed by intrinsic mechanical sensitivity of AN fibers and that this sensitivity underlies response properties that have not been well understood(Kiang, 1990). The use of presynaptic neurotransmission and postsynaptic mechanical transmission may be a cochlear feature that sculpts the extraordinary auditory response(Griesinger et al., 2005).

The findings from this report provide evidence for AN intrinsic mechanical sensitivity. If the AN’s presynaptic nerve endings directly sense sound-mediated OCM, then the distinctive classical role of the AN as primary neuron requires reevaluation. Moreover, the findings are in keeping with the emerging evidence that primary afferent neurons, such as visual retinal ganglion cells and DRG neurons, utilize melanopsin and mechano-sensitive channels, respectively, to modulate sensory processing(Coste et al., 2010; Hattar et al., 2002).

### Mechanical sensitivity of auditory neurons *in vitro*

The present study’s data show that SGN cell-body or dendrite displacement activates a membrane conductance with a reversal potential ∼0 mV, suggesting it is a nonspecific cationic current. The sensitivity of the displacement-induced current and membrane depolarization to GsMTx4-block(Bae et al., 2011) further indicates that SGNs express mechanically-sensitive channels. It was not the aim of this study to identify the candidate mechanically-sensitive pathway; however, several nonspecific cationic channels such as the transient receptor potential type 3 (TRPC3)(Phan et al., 2010), vanilloid I (TRPV1)(Zheng et al., 2003), and polycystine (TRPP2)(Takumida and Anniko, 2010) membrane proteins have been identified in SGN. Other putative mechanically-gated ion channels in SGNs can be derived from the library of single-neuron RNA-sequence analyses(Shrestha et al., 2018; Sun et al., 2018). These mechanically-gated channels include Piezo-1 and Piezo-2, identified in HCs and other cochlear cell-types(Beurg and Fettiplace, 2017; Corns and Marcotti, 2016; Shrestha et al., 2018). Null mutation of *Piezo*-2 in mice decreases the sensitivity of auditory brainstem responses by ∼20 dB. However, it is unlikely that Piezo-2-mediated currents in HCs account for the auditory phenotype seen in the null-mutant mouse. The Piezo-2-mediated current in HC declines during development and may be functionally insignificant after hearing(Beurg and Fettiplace, 2017). The identity of the well-studied, pore-forming mechanically-gated channel in HCs remains unknown(Giese et al., 2017; Qiu and Muller, 2018). If SGNs utilize a distinct mechanically-gated channel, a search for the gene and protein may include using emerging single-cell transcriptomic and phenotypic analyses of mouse models, which awaits future studies.

Some conditions of combined (current + displacement) stimulation (**Fig. 3, S10**) and even of displacement by itself (**S1, S2, S6**) caused suppression of spike rate of SGNs. While the data presented here cannot directly address the underlying mechanism for how two inwardly-directed currents (EPSC and I_MA_) yield suppressive effects, reasonable explanations can be considered. Recent reports have shown that besides voltage-dependence, outward K^+^ channels such as K_v_1.1/1.2 and Kv7 are modulated by mechanical stimuli(Hao et al., 2013; Perez-Flores et al., 2020), whereby the channel’s voltage sensitivity is enhanced(Long et al., 2005). K_v_1 and K_v_7 channels are members of the cadre of outward K^+^ channels that dominates the type I afferent AN membrane(Lv et al., 2010; Mo et al., 2002; Wang et al., 2013). If the magnitude of outward K^+^ currents enhanced by mechanical displacement exceeds the mechanically-activated inward currents, the resulting outcome would be membrane repolarization, which would suppress AN activity, consistent with the current results.

The suppression of AN fibers’ spiking is a known phenomenon *in vivo* and has been observed under two conditions. First, a sound can suppress the response to another sound (“two-tone suppression”)(Sachs and Kiang, 1968): while this is partly grounded in cochlear mechanical vibration, it remains unclear whether other mechanisms contribute(Robles and Ruggero, 2001; Versteegh and van der Heijden, 2013). Second, suppression of spiking can occur even in response to single, low-frequency tones of increasing intensity. Such stimuli trigger responses with an increasing number of spikes that are phase-locked at a preferred stimulus phase, but at high sound levels (80-90 dB SPL) a set of associated changes abruptly occurs: firing rate drops over a narrow range of intensities (“Nelson’s notch”)(Kiang and Moxon, 1972; Liberman and Kiang, 1984), and this is accompanied by a change in phase-locking towards multiple preferred phases (peak-splitting). At still higher intensities, the response returns to a high spike rate and monophasic phase-locking, albeit at values that differ from those at lower intensities. One functional consequence is increased coding of envelope fluctuations at high sound levels(Joris and Yin, 1992). Peak-splitting suggests an interaction of two pathways with different growth functions, which sum at the level of the AN(Kiang, 1990; Kiang et al., 1986). We observed phenomenologically similar events *in vitro* when combining sinusoidal current injection and displacement at varying relative phases. Some combinations generated peak-splitting, accompanied by decreases in firing rate (**Fig. 3, S10**). These findings suggest that the mechanical sensitivity of SGNs should be considered a possible factor in the suppressive and peak-splitting phenomena observed *in vivo*.

Technical limitations restricted *in vitro* experiments to much lower frequencies (by about a factor of 10) than those at which peak splitting is typically studied *in vivo*. However, note that peak-splitting becomes more prevalent and occurs over a broader range of sound levels when the sound frequency is lowered to values similar to those used here *in vivo*(OshI_MA_ and Strelioff, 1983; Ronken, 1986; Ruggero and Rich, 1983). We surmise that mechanical displacement or deformation of SGN dendrites affects their response to sound over a broad range of frequencies, but with a bias towards low frequencies. Moreover, mechanical effects are bound to occur at a particular phase relationship relative to the synaptic drive on the same dendrite, and that relationship likely varies with stimulus frequency and possibly also with the cochlear longitudinal location. The relative magnitude and phase of synaptic versus mechanical stimulation would determine the phase and probability of spiking.

### Mutually interacting elements of neurotransmission and mechanical activation at the first auditory synapse

Besides the complexities in phase-locking that AN fibers show *in vivo* in response to simple tones, complexities in frequency tuning are observed as well: both sets of phenomena point to the existence of multiple components driving AN responses(Kiang, 1990; Kiang et al., 1986; Kiang and Moxon, 1972; Liberman and Kiang, 1984). In the cat, the species in which this was first and most extensively described, a characterization in terms of two components was proposed, with one component dominating at low sound levels and a second component at high sound levels(Kiang, 1990; Liberman and Kiang, 1984). Similar phenomena, particularly regarding phase-locking, have been reported in other species but were not always restricted to high sound levels. The source of these components is controversial but has been sought at the level of cochlear mechanics or hair cells, not at the AN(Cai and Geisler, 1996; Cody and Mountain, 1989; Dallos, 1985; Heil and Peterson, 2019; Kiang, 1990; Liberman and Kiang, 1984; Ruggero and Rich, 1983; Ruggero et al., 1986)^72^.

We hypothesized that intrinsic mechanical sensitivity contributes to the high-threshold, “tail” region of tuning curves, which is less vulnerable to a range of cochlear manipulations than the tip(Kiang et al., 1986; Liberman and Kiang, 1984). These tails may be associated with clinically relevant phenomena, such as recruitment of middle ear reflexes and abnormal growth of loudness after acoustic trauma(Liberman and Kiang, 1984). Delivery of synaptic blockers diminished or abolished spontaneous activity and the tip region of the tuning curve. The observation that AN responses persisted at high sound levels is in keeping with the prediction that a second mode, direct mechanical activation of AN fibers, operates and dominates high-threshold segments of the tuning curve (**Fig. 4, S11**).

Suppose neurotransmitter release-mediated EPSCs are the sole driver of AN responses as suggested(Fuchs et al., 2003), and suppose the mechanisms underlying the peculiarities in frequency tuning precede the synapse between IHC and AN fiber. In that case, synaptic blockers should attenuate AN activity at all intensities and should not differentially affect the presence of multiple components. This was not what we observed: while low-threshold responses were abolished, spikes could still be elicited at high intensities (**Fig. 4a-f**), even in conditions where cochlear mechanical sensitivity and mass receptor potentials were normal (**Fig. 4g**). It is conceivable that with strong depolarization, and by extension, high sound levels, the copious release of glutamate at the synapse may render a competitive antagonist, such as NBQX, ineffective(Sheardown et al., 1990). Arguing against this explanation is that similar results were obtained with an L-type Ca^2+^ channel blocker (**S11**). NBQX and Ca^2+^ channel blockers’ differential effects on the two components of AN tuning curves (**Fig. 4, S11**) argue against a single-element (neurotransmitter) mechanism(Fuchs et al., 2003; Glowatzki and Fuchs, 2002). The results are consistent with two mutually interacting mechanisms: neurotransmission and direct mechanical activation.

It may be argued that dual neurotransmitter release properties perhaps explain the two components of AN tuning curves. Under certain conditions, IHC synaptic release mechanisms may consist of two distinct modes(Grant et al., 2010), which are observed at steady-state and contribute towards spontaneous APs in afferent fibers. If the two modes of synaptic transmission represent the two components of the AN tuning curve, again, NBQX and Ca^2+^ channel blockers are expected to affect both components(Glowatzki and Fuchs, 2002; Grant et al., 2010), which is not what we observed: NBQX and Ca^2+^ channel blockers suppressed the sharp-frequency tip at the CF, with a remaining response showing very coarse tuning (**Fig. 4, S11**). Thus, it is unlikely that the biphasic modes of transmitter release mechanisms, or, for that matter, any synaptic or presynaptic mechanisms, account for the differential effect of blockers on the low- and high-threshold components of the AN tuning curve. We propose, therefore, that the element of the tuning curve that remains impervious to the synaptic transmission block is driven mainly by mechanical activation of the AN.

Small molecule blockers of mechanically-gated ion channels, such as Gd^3+^, are nonspecific in action(Ranade et al., 2015) due to their effects on Ca^2+^ and K^+^ channels and synaptic transmission and cannot be used to suppress SGN mechanical-sensitivity selectively. Moreover, diffusional constraints for large molecular-weight channel blockers (e.g., GsMTx4) also preclude *in vivo* selective blocking of SGN mechanical-sensitivity.

### Mechanical sensitivity of auditory neurons *in vivo*

The *in vivo* approach faced experimental constraints. Unavoidably, blind recording from the nerve trunk biases against neurons lacking spontaneous activity and low-threshold responses. However, the most severe experimental deficiency is the diffusion of blockers through long and narrow membrane-bound spaces to reach the IHCs and free dendritic endings of SGNs. The observation that apical neurons tuned to low frequencies were invariably less affected than neurons innervating more basal parts of the cochlea suggests that the drugs delivered through the round window did not always reach the HC-AN synapse at more apical locations. Diffusional limits also prevented the use of larger molecules, such as GsMTx4.

The *in vitro* results suggest that synaptic and mechanical effects do not merely superimpose but interact in a nonlinear fashion. For example, if one modality (current injection or mechanical deformation) is subthreshold, it can be made suprathreshold by adding the other modality (**Fig. 2**). This complicates the interpretation of the *in vivo* experiments – a complete absence of spiking after blocking synaptic transmission does not rule out a role for intrinsic mechanical sensitivity, and a partial effect of synaptic blockers on spiking does not prove such a role. Therefore, the *in vivo* experiments do not conclusively demonstrate that responses at high sound levels depend on intrinsic mechanical sensitivity, but they are consistent with that proposal.

Although the thresholds of substrate displacement needed to trigger spiking *in vitro* are within the range of displacements of cochlear structures measured *in vivo* at high sound levels(Cooper and Rhode, 1995; He et al., 2018; Lee et al., 2016a), it is at present unclear what the displacement amplitudes are *in vivo* at the unmyelinated dendritic terminals of the AN. Recent measurements with optical coherence tomography reveal a microstructure of displacements, with the largest displacements occurring at low frequencies at locations away from the basilar membrane (Cooper et al., 2018; He et al., 2018; Lee et al., 2015). Undoubtedly, the substrate-displacement stimuli used in the present study are only a coarse approximation of the *in vivo* OCM. The relevant physical stimulus is possibly in small pressure differences, rather than displacements, that follow the acoustic waveform between the OC and the modiolar compartment and cause deformations of dendrites of AN fibers(Karavitaki and Mountain, 2007) and trigger intrinsic mechanical effects. Sensitivity of AN fibers to displacements or pressure differences suggests new treatment strategies for the hearing impaired.

## Acknowledgments

We thank members of our laboratories for comments on this manuscript. We thank Dr. Frederick Sachs for his constructive comments and suggestions on the first draft of the script. Grants to ENY supported this work from the National Institutes of Health (DC016099, DC015252, DC015135, AG060504, AG051443). PXJ was supported by grants from KU Leuven (BOF, OT-14-118) and Fonds Wetenschappelijk Onderzoek (Research Foundation Flanders) G0B2917N.

Supplementary Information is available in the online version of the paper.

## Author Contributions

ENY and PXJ designed the research; MCPF, EV, PXJ, and ENY analyzed data and wrote the manuscript; MCPF, JHL, PXJ, EV, HJK, and ENY performed the research. All authors read and approved the final manuscript.

## Author Information

Correspondence should be addressed to ENY, Program in Communication Science, Department of Physiology and Cell Biology, University of Nevada School of Medicine, Reno, NV.

## Methods

All *in vivo* experiments were performed under a protocol approved by the University of Leuven’s animal ethics committee complying with the European Communities Council Directive (86/609/EEC). At the University of Nevada, Reno (UNR), experiments were performed according to the guidelines of the Institutional Animal Care and Use Committee of UNR. *In vitro* experiments were performed using mice and chinchilla. Equal numbers of adult male and female mice and chinchillas were used (when odd numbers are reported, the females outnumbered males). *In vivo* experiments were performed on adult wild-type chinchilla (*Chinchilla lanigera*) of both sexes, free from a middle ear infection and weighing between 200-700g.

### Cell culture

Spiral ganglion neurons (SGNs) were isolated from five-to-eight-week-old C57BL/6J mice (Jackson Laboratory) and one-to-two-month-old (200-300g) chinchilla as described previously(Lee et al., 2016b). The apical and basal cochlea SGNs were dissociated using a combination of enzymatic and mechanical procedures. The neurons were maintained in culture for two to five days. Animals were euthanized, and the temporal bones were removed in a solution containing Minimum Essential Medium with HBSS (Invitrogen), 0.2 g/L kynurenic acid, 10 mM MgCl_2_, 2% fetal bovine serum (FBS; v/v), and 6 g/L glucose. The central spiral ganglion tissue was dissected out and split into three equal segments: apical, middle, and basal, across the modiolar axis, as described previously^24^. The middle turn was discarded, and the apical and basal tissues were digested separately in an enzyme mixture containing 1 mg/ml collagenase type I and 1 mg/ml DNase at 37°C for 20 min. We performed a series of gentle trituration and centrifugation in 0.45 M sucrose. The cell pellets were reconstituted in 900 μl of culture medium (Neurobasal-A, supplemented with 2% B27 (v/v), 0.5 mM L-glutamine, and 100 U/ml penicillin; Invitrogen), and filtered through a 40 μm cell strainer for cell culture and electrophysiological experiments. For adequate voltage-clamp and satisfactory electrophysiological experiments, we cultured SGNs for ∼24–48 hours to allow Schwann cells’ detachment from neuronal membrane surfaces.

Chinchilla were purchased from Moulton Chinchilla Ranch (Rochester, MN). Chinchilla SGNs were isolated using a protocol similar to that employed for mice. The tendency for neuronal culture from the chinchilla to profusely generate glial cells was high. We inhibited glial-cell proliferation, using 20 μM cytosine arabinoside (AraC; Sigma)(Schwieger et al., 2016) in the culture solution. Electrophysiological experiments were performed at room temperature (RT; 21-22°C). Reagents were obtained from Sigma Aldrich unless otherwise specified.

### Electrophysiology

Experiments were performed in standard whole-cell recording mode using an Axopatch 200B amplifier (Axon Instruments). For voltage-clamp recordings patch pipettes had resistance of 2-3 MΩ when filled with an internal solution consisting of (in mM): 70 CsCl, 55 NMGCl, 10 HEPES, 10 EGTA, 1 CaCl_2_, 1 MgCl_2_, 5 MgATP, and 0.5 Na2GTP (pH adjusted to 7.3 with CsOH). The extracellular solution consisted of (in mM): 130 NaCl, 3 KCl, 1 MgCl_2_, 10 HEPES, 2.5 CaCl_2_, 10 glucose and 2 CsCl (pH was adjusted to 7.3 using NaOH). For current-clamp recordings, pipettes were filled with a solution consisting of (in mM) 134 KCl, 10 HEPES, 10 EGTA, 1 CaCl_2_, 1 MgCl_2_, and 5 MgATP and 0.5 Na2GTP (pH 7.3 with KOH). Currents were sampled at 20-50 kHz and filtered at 2-5 kHz. Voltage offsets introduced by liquid junction potentials (2.2 ± 1.2 mV (n = 45)), were not corrected. Leak currents before mechanical stimulations were subtracted offline from the current traces and were <20 pA. Recordings with leak currents greater than 20 pA were discarded. A stock solution of 10 mM GsMTx4 (CSBio; Menlo Park, CA) was prepared in water.

### Mechanical stimulation

Mechanical stimulation was achieved using a fire-polished and sylgard-coated glass pipette (tip diameter ∼1 μm), positioned at ∼180° to the recording electrode. The probe’s movement toward the cell was driven by a piezoelectric crystal micro stage (E660 LVPZT Controller/Amplifier; Physik Instruments). The stimulating probe was typically positioned close to the cell body without any visible membrane deformation. The stimulation probe had a velocity < 20 μm ms^-1^ during the ramp segment of the command for forwarding motion, and the stimulus was applied for a duration, as stated in each experiment. We assessed for mechanical sensitivity using a series of mechanical steps in ∼0.14 μm increments applied every 10 to 20 s, which allowed for the full recovery of mechanosensitive currents between steps. I_MA_ were recorded at a holding potential of -70 mV. For instantaneous I-V relationship recordings, voltage steps were applied 8 s before the mechanical stimulation from holding potentials ranging from -90 to 90 mV.

SGNs were cultured on a polydimethylsiloxane (PDMS) substrate treated with poly-D-lysine (0.5 mg/ml) and laminin (10 mg/ml) to test for the mechanosensitivity of nerve endings. A single neurite can be stretched by substrate indentation on this platform without contacting the neurite (**Fig. 1b** **inset**). The whole-cell patch-clamp recording was used to examine the electrical response to neurite stretching conducted with our direct approach of indentation of a PDMS substrate at a location adjacent to the neurite with a pipette. To study the neurons’ firing response to time variation with simultaneous sine wave current and mechanical stimulation, the mechanical stimulus phase was shifted in 45° steps from 0 to 315° relative to the current. Positive voltage to the actuator corresponds to downward displacement. Technical limitations restricted *in vitro* experiments sinusoidal mechanical stimulation to ∼100 Hz and current injection to ∼1000 Hz.

### Cryosection

The temporal bones were removed and fixed in 4% paraformaldehyde in phosphate-buffered saline (PBS) for 1.5 hr at 4°C. The temporal bones were decalcified by incubation in 10% EDTA at 4°C for 3-5 days. The EDTA solution was changed daily. The bones were then embedded in OCT compound for cryostat sectioning. The sections of 10 μm thickness were washed in PBS, and nonspecific binding was blocked with 1% bovine serum albumin (BSA) and 10% goat serum in PBS plus 0.1% Triton X-100 (PBST) for 1 hr. The primary antibodies, chicken anti-Tuj1 (Abcam), mouse anti-myelin basic protein (Abcam), and rabbit anti-Myo7a (Proteus Biosciences, Inc.), were incubated overnight at 4°C. After the incubation with the primary antibodies, the slides were washed three times with PBST and incubated with secondary antibodies for 1.5 hours at room temperature in the dark. We used Alexa Fluor 647-conjugated goat anti-mouse and Cy3-conjugated goat anti-chicken, and Alexa Fluor 488-conjugated goat anti-rabbit in a dilution of 1:500. The slides were then examined under a confocal microscope (LSM 510, Zeiss).

### Data analysis

Data analyses were performed offline using pClamp8 (Axon Instruments) and Origin software (Microcal Software, Northampton, MA). Statistical analysis was performed using a paired or unpaired *t-test*, with significance taken at *p* < 0.05. The peak I_MA_ for each step displacement was expressed in channel-open probability (*P*_o_) and plotted against displacement (X). The relationship was fitted with a one-state Boltzmann equation *P_o_* = 1/[1 + e*^z^*^(^*^X^* − *^X^0.5*^)/^(*kT)*] to obtain channel gating force, *z*, and the displacement at 50% open probability (*X_0.5_*), and *T* is temperature. For a two-state Boltzmann function *P*_o_ = 1/([1 + e*^za^*^(^*^X^* − *^X^0.5a*^)/^(*kT)*]+ 1/[1 + e*^zb2^*^(^*^X^* − *^X^0.5b*/(*kT)*]). The dose-response relationships were described with a logistic function; (I_i_ – I_f_)/(1 + ([C]/[C_0.5_])*^p^* + I_f_. I_i_ and I_f_ are the initial and final magnitudes, [C] is the drug’s concentration with [IC_0.5_] its half-blocking concentration, and *p* is the Hill coefficient. The decay phases of the I_MA_ were fitted by a bi-exponential decay function of the form: *y*(*t*) =*A*_1_ * exp(−*t*/τ_1_) + *A_2_* * exp(−*t*/τ_2_) + *A*_ss_, where *t* is time, τ is the time constant of decay of IMA, *A*_1_ and *A*_2_ are the amplitudes of the decaying current components, and *A*ss is the amplitude of the steady-state, non-inactivating component of the total IMA. The strength of phase locking was quantified as vector strength (VS), which is the ratio of the period histogram’s fundamental frequency component to the average firing rate^36^. Also known as the synchronization index, the VS varies from VS = 0 for a flat histogram with no phase locking to VS = 1 for a histogram indicating perfect phase locking. Statistical significance of the VS was assessed by Rayleigh statistics(Mardia, 1972), using a criterion of *p* < 0.05; values of VS failing this criterion were discarded. Data are presented as mean ± SD (standard deviation).

### Surgical preparation for *in vivo* experiments

In chinchillas, anesthesia was initiated by intramuscular (im) injection of a ketamine-xylazine mixture and was maintained with ketamine and diazepam, titrated according to vital signs and reflexes. In cats, induction was with a 1:3 mixture of ketamine and acepromazine and maintenance with intravenous infusion of pentobarbital. The animal was placed in a double-walled sound-proof room on a feedback-controlled heating pad. A tracheotomy was performed, and the respiration rate and end-tidal CO2 were continuously monitored. The acoustic system was placed in the external auditory meatus and calibrated with a probe microphone. The AN was accessed via a traditional posterior fossa approach in which a small portion of the lateral cerebellum was aspirated. The tympanic bulla on the recording side was opened to visualize the cochlear round window for cochlear potential measurements and blockers’ administration. Details of our procedures are available elsewhere(Bremen and Joris, 2013; Louage et al., 2006).

### Acoustic stimulation and recording

Acoustic stimulation and acquisition of signals utilized custom software to control digital hardware (RX6, Tucker-Davis Technologies, Alachua, FL). Stimuli were compensated for the transfer function of the acoustic system. Acoustic stimuli were delivered through dynamic phones. For CM and DPOAE recordings, analog signals were recorded (RX6) and analyzed off-line (MATLAB, MathWorks, Natick, MA). CM recordings were obtained with a silver ball electrode near the RW. The reference and ground wire electrodes from the differential amplifier were placed in the skin next to the ear canal and in the neck’s nape, respectively. The signal was amplified with a differential amplifier (RS560, Stanford Research Systems), recorded (RX8, Tucker-Davis Technologies, Alachua, FL), stored on a computer, and processed with custom software in MATLAB. The CM was obtained for different stimulus frequencies (2,3,4,6,8,10,12 kHz) and was spectrally calculated from the difference of the evoked responses (divided by 2) to alternating pure tones. DPOAE responses were recorded and stored using the same microphone and acquisition system as for the acoustic calibration. The primaries (F1 and F2, duration 700 ms, repetition interval 800 ms) were generated with separate acoustic actuators to minimize actuator distortion products. The frequency ratio (F2/F1) and amplitude levels (L1, L2) of the primaries were fixed to a ratio of 1.21 and 65 (L1) and 55 (L2) dB SPL, respectively. DPOAEs were quantified as the sound pressure of the returning 2F1-F2 cubic difference distortion product.

Single fiber recordings were obtained with micropipettes filled with 2M KCl (impedance ∼40-80 MΩ), mounted in a hydraulic micro-drive, and placed above the nerve trunk under visual control. Zero depth was marked at initial contact. If necessary, warm agar (2%) was poured on the AN to reduce pulsations. The neural signal was recorded and displayed using routine methods. An intracellular amplifier (Dagan BVC-700A) allowed monitoring of DC-shifts signaling axonal penetration and enabled the use of current pulses to improve the recording.

Further amplification and filtering (∼ 100 Hz-3 kHz) preceded conversion of spikes to standard pulses with a custom peak-detector (1 µs resolution) and internal RX6 timer. A search stimulus consisting of 60 dB SPL, 200ms tone pips stepped in frequency was used while the microelectrode was advanced in 1-µm steps until large monopolar action potentials were obtained. After the blocker administration, the search stimulus’s sound level was gradually increased, and the duration of the tone pips reduced to minimize cochlear trauma. For each isolated AN-fiber, the spontaneous rate (SR: estimated over a 15-second interval) was measured, and a threshold tuning curve was obtained with a tracking program using short (30 ms) tone bursts (repetition intervals 100 ms; rise-fall time 2.5 ms). We extracted the characteristic frequency (CF) as the frequency of the lowest threshold from the threshold tuning curve.

---

**Figure S1.**
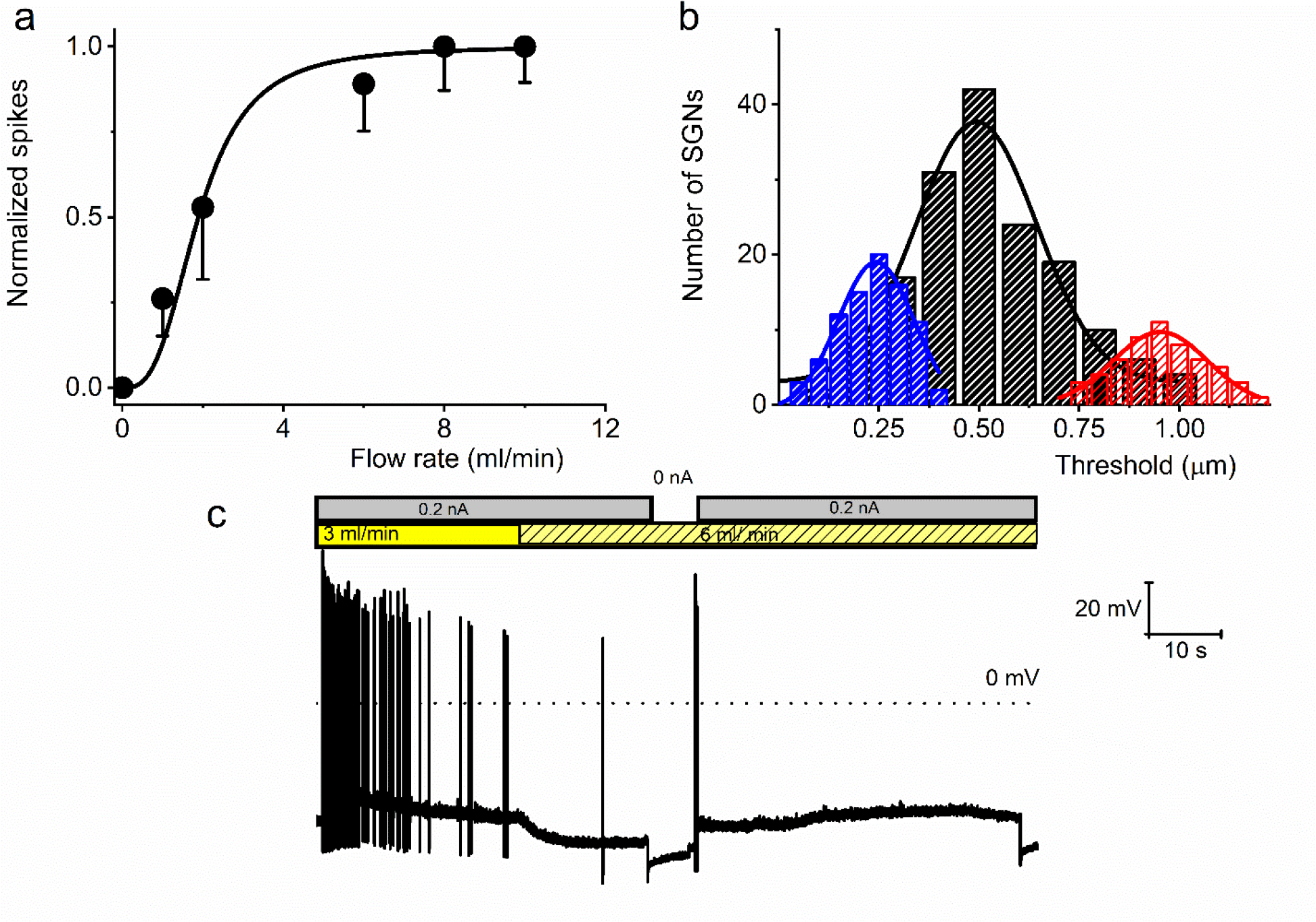
Effect of bath solution flow rate on the SGN firing rate. **a**, Summary data from four eight-week-old SGNs showing changes in normalized spike rate at different bath solution flow rates. The normalized spike rate saturated at ∼8 ml/min. **b**, Cumulative binomial distribution of the mechanical threshold (mean cell-body displacement in μm) to elicit APs in mechanically-sensitive SGNs. Three distinct peaks were observed: 0.24 ± 0.01, n = 85 (blue); 0.51 ± 0.01, n = 164 (black); and 0.95 ± 0.02, n = 56 (red). **c**, A membrane response of an eight-week-old apical SGN at a current injection of 0.2 nA and perfusion flow rates of 3 and 6 ml/min. The shift from slow (3 ml/min) to fast (6 ml/min) rate caused membrane hyperpolarization and reduced spike activity.

**Figure S2.**
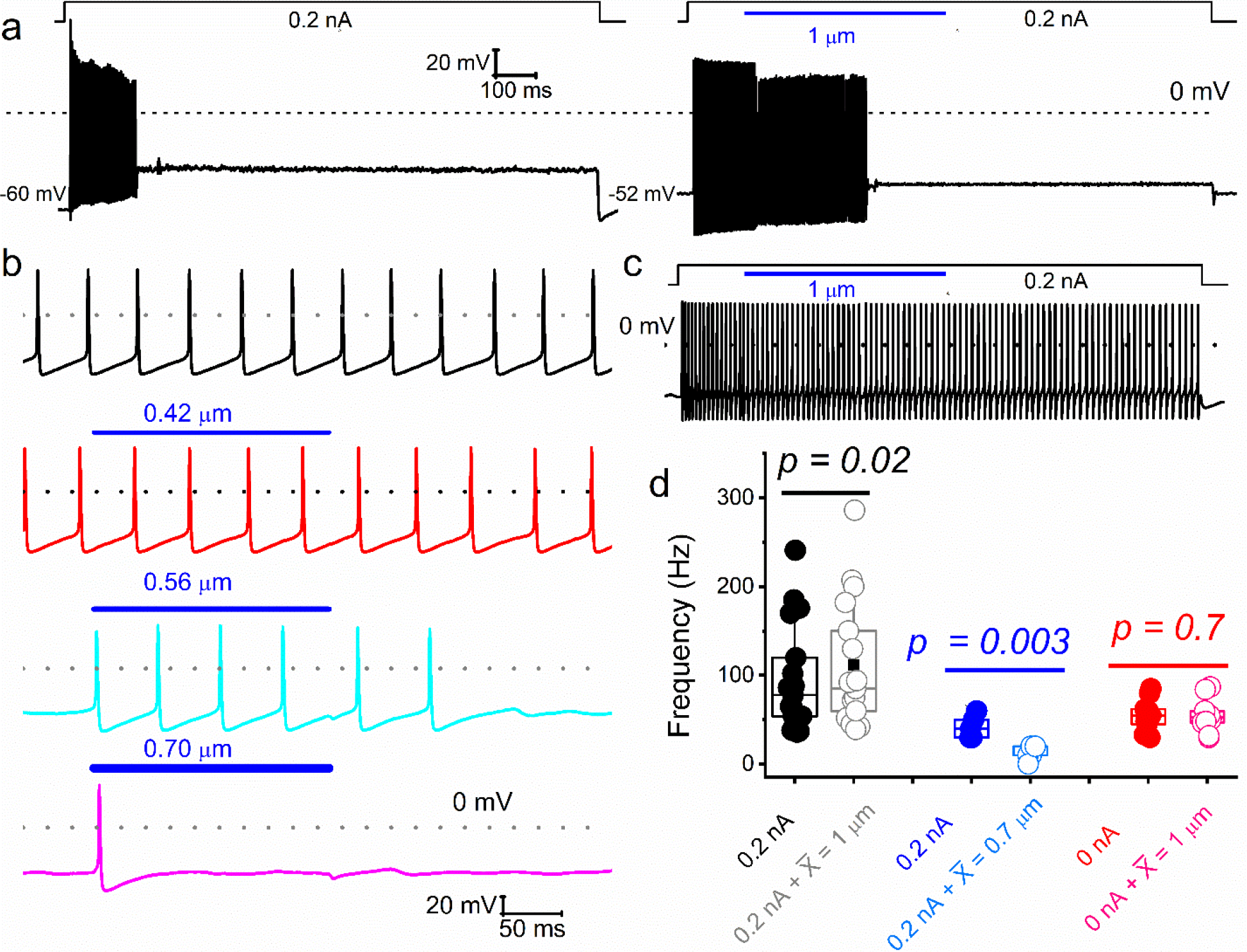
Varied responses to combined mechanical and electrical stimulation of SGNs. Representative current-clamp responses evoked by current injection and combined mechanical displacement (blue) were applied in soma-displacement (1 μm). **a**, Shown is an example of a basal SGN response injected with a 0.2 nA current alone (left panel) to invoke firing responses, which were subsided despite maintained current-injection. When both stimuli are applied simultaneously (right panel), the firing rate increased. Note that the spike activity halted before ending the mechanical displacement. V_rest_ shifted by ∼8 mV in depolarizing direction, in the right trace, which may account for the spike amplitude reduction. The firing rate increased in many (∼38%; 12 out of 32 SGNs) adapting neurons when the current injection was combined with mechanical displacement. **b**, In spontaneously active SGNs, gradual increase (0 to 0.7 μm; see lower panels) in dendritic substrate displacement resulted in reduced spike rate (31%; 10 out of 32 SGNs). **c**, For this non-adapting basal neuron, the firing rate remained unchanged (∼31%; 10 out of 32 SGNs). **d**, Summary data, showing the response properties of SGNs to current alone and combined current and mechanical displacement (indicated). Data are grouped into SGN subgroups in which paired current and mechanical displacement resulted in increased (black symbols), reduced (blue), and unaltered (red) spike frequencies.

**Figure S3.**
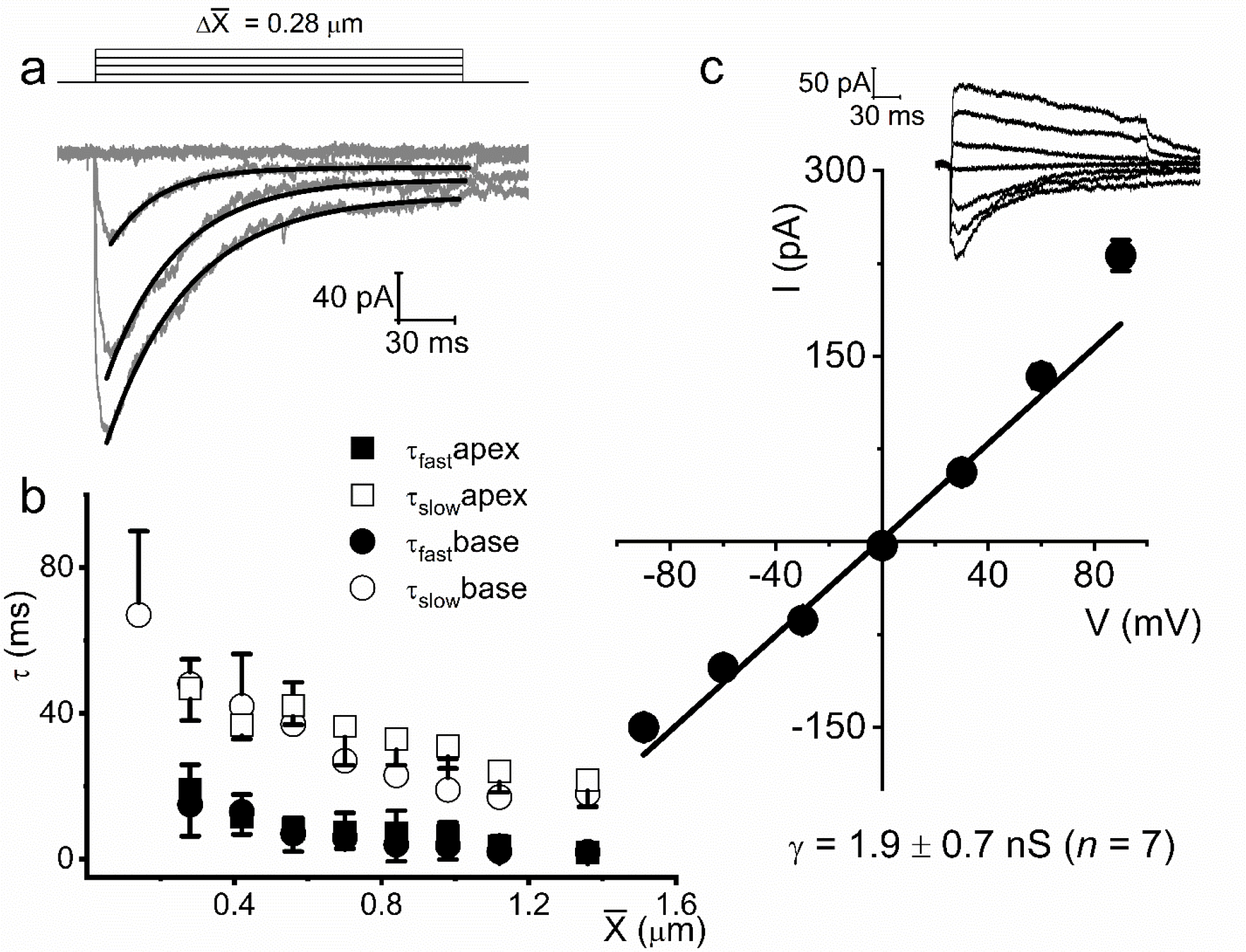
Displacement-clamp current recordings from eight-week-old adult SGNs. **a**, Current traces (dark gray) in response to ∼215 ms mechanical stimulus steps of ∼0.28 μm increments in SGNs from the basal cochlear turn in the mouse. The holding potential was -70 mV. The solid black lines represent the double exponential fits for the current decay kinetics. **b**, Summary data of the decay time constant (τ) as a function of displacement for (X) for currents recorded from SGNs from apical (*n* = 17) and basal (*n* = 15) cochleae. **c**, Shown is the average I-V relationship of MA currents evoked at different membrane potentials (-90 to 90 mV), using ∼0.4 μm displacement. The regression line indicates an MA current reversal potential (E_MA_) of - 2.7 ± 2.1 mV (*n* = 13). The inset shows an example of representative MA current (I_MA_) traces with membrane voltages stepped in 30 mV increments, from -90 to 90 mV.

**Figure S4.**
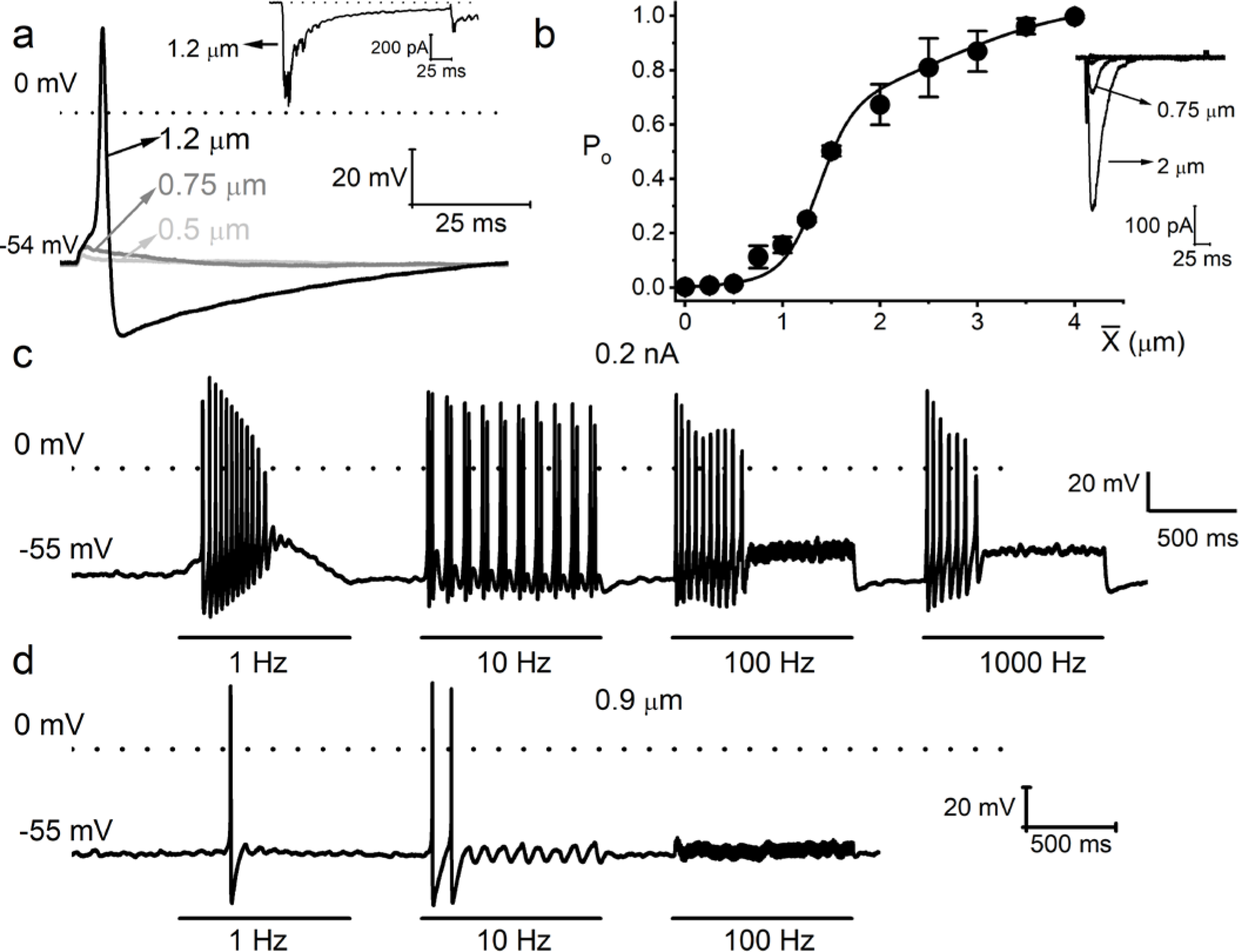
Mechanical sensitivity of mouse vestibular ganglion neurons. **a**, Response properties of a six-week-old mouse vestibular neuron to square pulse mechanical displacements. The magnitudes of the displacement at the soma are indicated. The dotted line denotes 0 mV, and the resting membrane potential (V_rest_) is indicated. The inset shows the inward current generated from the same neuron in voltage-clamp mode at -70 mV holding voltage. **b**, Summary data of displacement-response relationship of I_MA_ represented as channel open probability (P_o_) as a function of displacement, fitted by a two-state Boltzmann function. The one-half-maximum displacements (X_0.5_) were 1.37 ± 0.10 μm and 2.60 ± 0.97 (n = 7). Inset shows example traces elicited in a mouse vestibular neuron. **c**, Eight-week-old vestibular neurons were injected with sinusoidal current with 0.2 nA peak-to-peak amplitude and varying frequencies (in Hz, 1, 10, 100, and 1000). **d**, Mechanical displacements were applied to the same cell using similar oscillatory frequencies with a peak-to-peak amplitude of ∼0.9 μm to elicit APs. Only rates at 1 and 10 Hz sufficed to elicit spikes.

**Figure S5.**
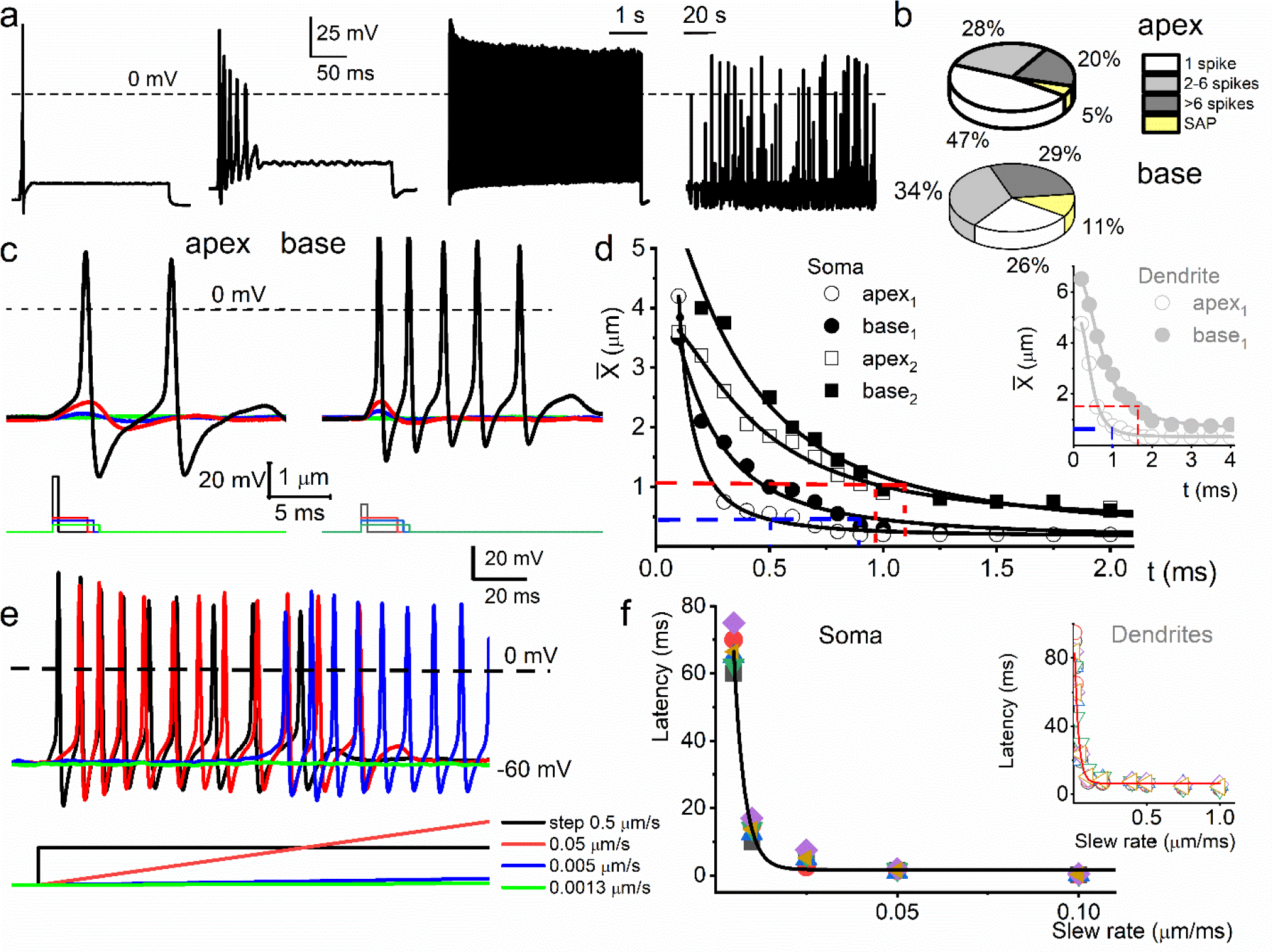
Response properties of adult mouse SGNs to current injection and mechanical stimulation. **a**, Different response properties from basal SGNs to 0.2 nA injection, from left to right panel: fast adapting (1 spike/stimuli), moderately adapting (2-6 spikes/stimuli), slow adapting (>6 spikes/stimuli), and neurons that fire without current injection (spontaneously firing; spontaneous action potential (SAP). **b**, Pie charts illustrate the proportions between the different types of SGN responses (∼five-to-six-week-old mice; 0.2 nA injection) for the apex (upper panel; n = 72) and the base (lower panel; n = 68). **c**, Membrane voltage responses to the cell-body displacement, from apical (left panel) and basal (right panel) cochlear neurons. The amplitude and duration of soma displacement were in the form of rectangular steps, as indicated. The amplitude and duration for the left panel (apical SGN) were as follows (in μm and ms), 2 and 0.5 (black); 0.5 and 3 (red); 0.4 and 3.5 (blue); 0.25 and 4 (green). The amplitude and duration for the right panel (basal SGN) were as follows (in μm and ms), 1 and 0.5 (black); 0.5 and 3 (red); 0.4 and 3.5 (blue); 0.25 and 4 (green) **d**, A plot of mean rheobase using soma displacement of apical and basal neurons and their respective chronaxie indicated in red and blue dashed lines. Mean rheobase displacement of dendritic substrate for SGNs from the cochlea’s apex and the base is shown in the inset. **e**, Family of APs evoked using step displacement (0.5 μm) and varying slew rates, indicated by color codes. The differences in latency of AP initiation are visible. **f**, The plot shows the relation between AP latency and the slew rate of mechanical displacement (*n* = 5). AP latency using the indirect movement of dendrites through substrate displacement at varying slew rates for 4 SGNs is plotted as inset.

**Figure S6.**
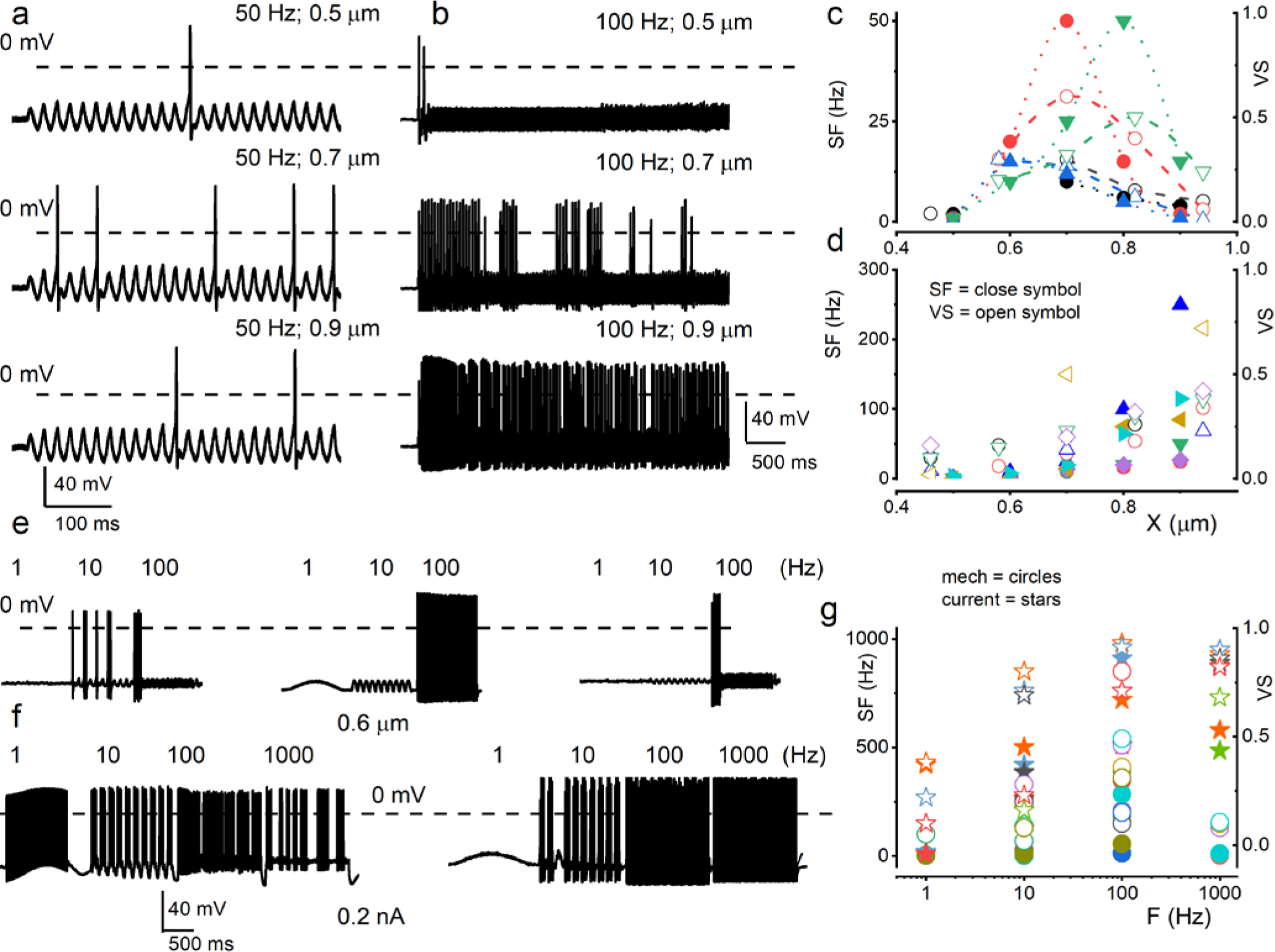
Mouse SGN response properties to sinusoidal mechanical and current stimulation. **a**, Characteristic apical SGN stimulated with 0.5-μm mechanical-displacement at 50 Hz. Subsequent traces below show the responses with 0.7- and 0.9-μm sine wave displacements at 50 Hz. **b**, Another apical SGN presented with 0.5-mm mechanical-displacement at 100-Hz and increasing stimulus amplitudes as indicated. The data represent SGNs that elicit increasing APs frequency with increasing mechanical displacement amplitude (**b**) and ones that show peak firing at specific displacement amplitude (**a**). **c-d**, Summary data of the two sets of SGNs, showing the relationship between displacement amplitude, spike frequency (SF; in solid symbols) and vector strength (VS; in open symbols). Each symbol represents a different SGN (**c**, n = 4; **d**, n = 6). **e**, Exemplary three different basal SGN stimulated with constant displacement (0.6 μm) at 1, 10, and 100 Hz. When presented with a constant displacement, neurons appeared to respond at a narrow frequency range (1-100 Hz), contrasting to current injection (1-1000 Hz) (**f**). Summary data show the relationship between displacement or current frequency, spike frequency (SF; in solid circles) and vector strength (VS; in open circles). Each symbol represents a different SGN (n = 6). Data from the current injection are represented with star symbols (n = 5).

**Figure S7.**
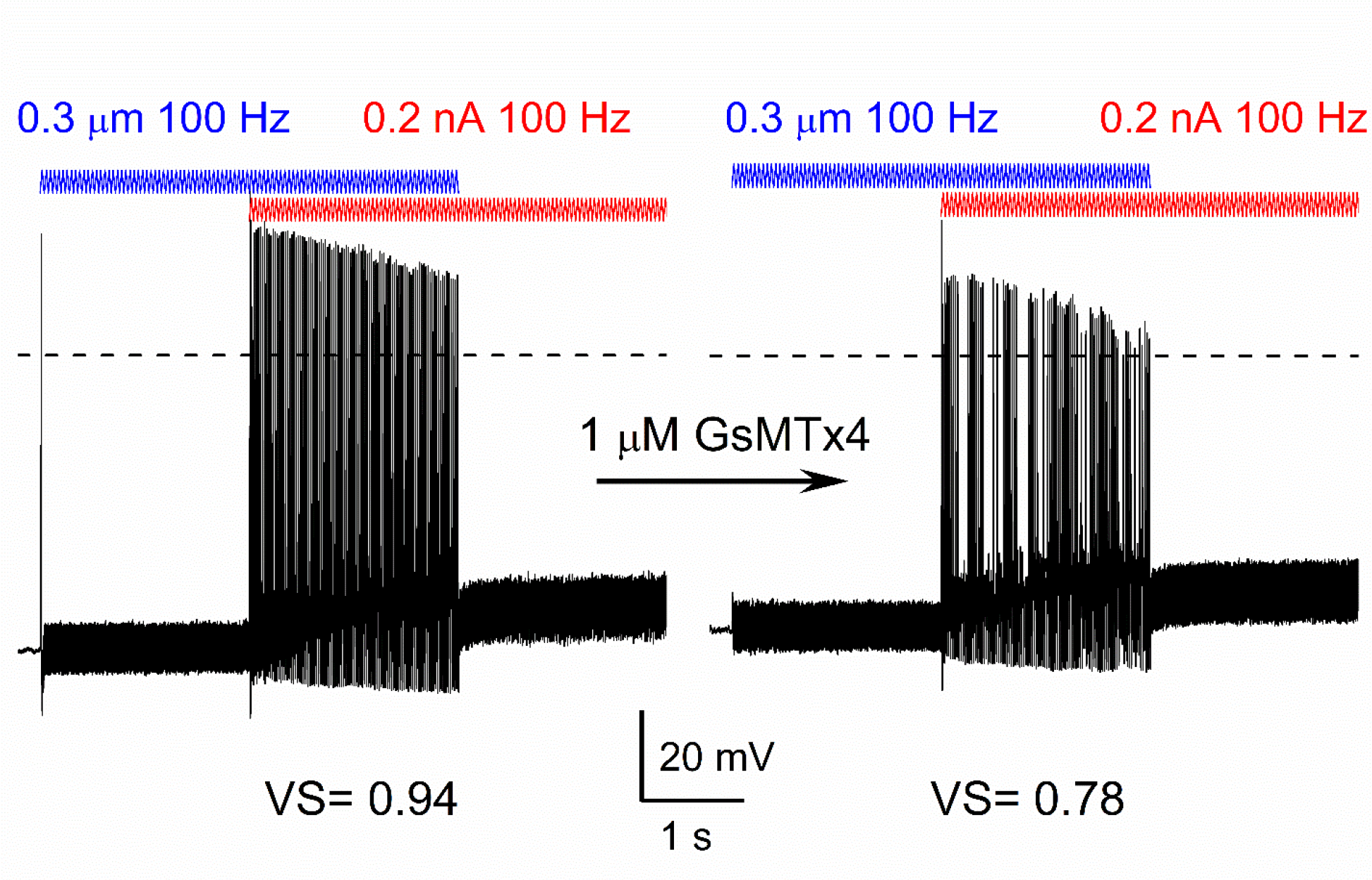
GsMTx4 reduced facilitation and spike timing after dual current injection and mechanical stimulation. Bath perfusion of GsMTx4 (1 μM), decreased facilitation, and spike timing resulting in reduced vector strength (VS) of the response to the simultaneous application of 100 Hz mechanical displacement (0.3 μm, blue) and current injection (0.2 nA, red) in an eight-week-old SGN.

**Figure S8.**
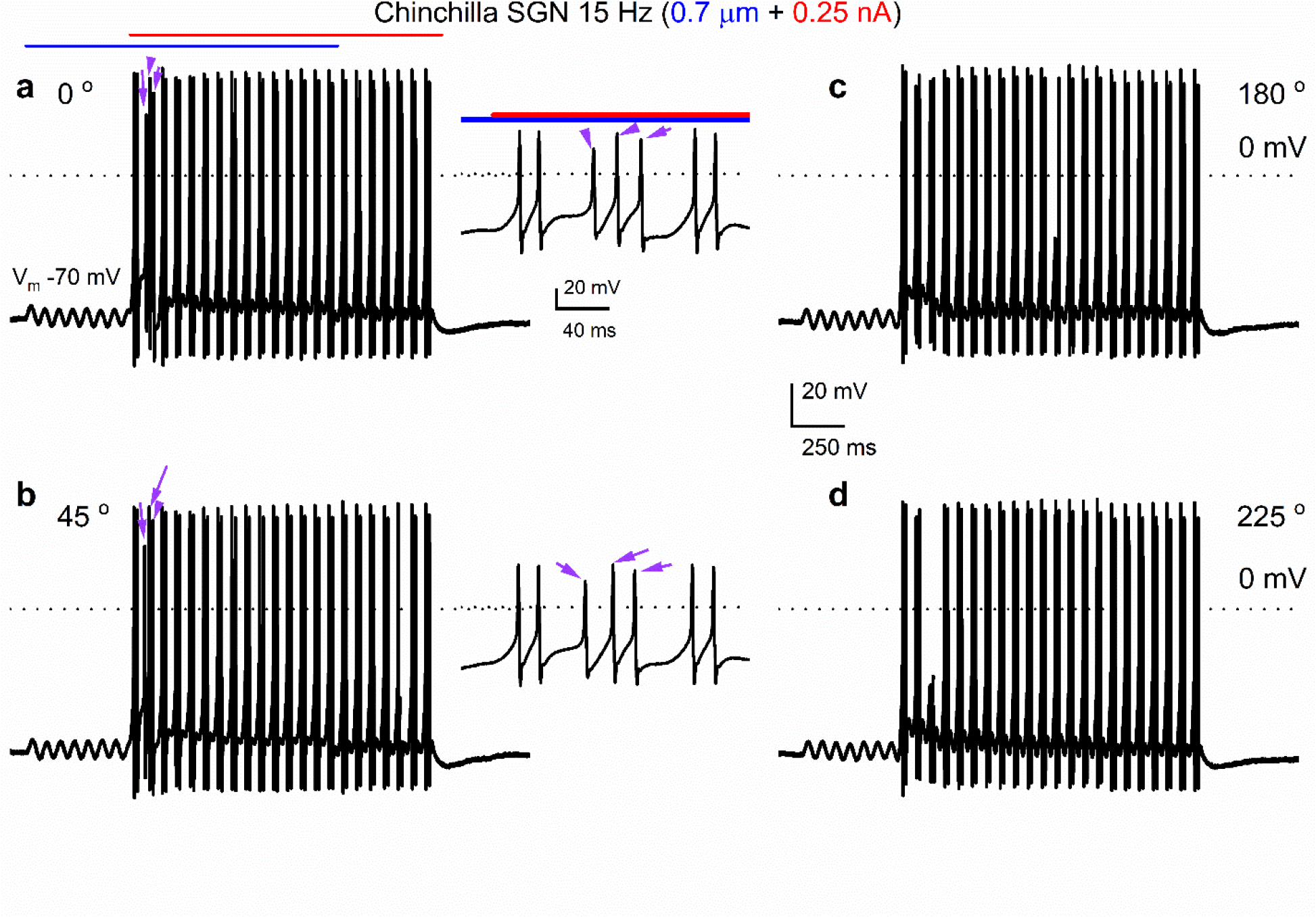
Examples of combined displacement and current injection yield more than two action potentials per cycle in chinchilla SGNs. Shown are voltage responses of chinchilla SGNs evoked with sinusoidal (15 Hz) mechanical displacement (0.7 μm, in blue) combined with sinusoidal (15 Hz) current injection (0.25 nA, in red) at four different relative phase angles (a-d, 0°, 45°, 180°, 225°, respectively). For most cycles, two action potentials were elicited per cycle, and this was the case for all four relative phase angles tested. However, as seen in a and b, the second cycle yielded three action potentials (purple arrows) for the 0° and 45° relative phased angles (insets).

**Figure S9.**
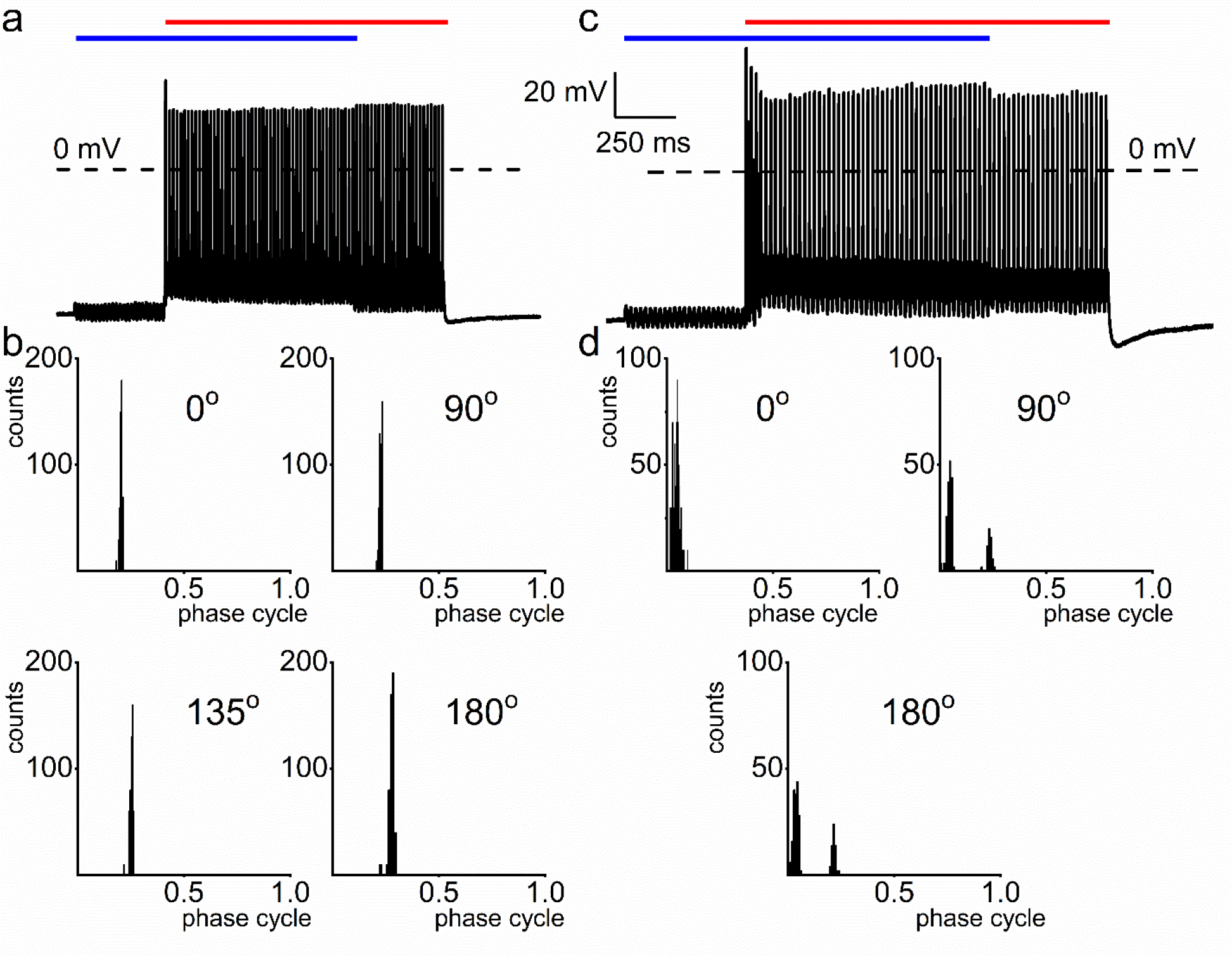
Mechanical sensitivity of chinchilla SGNs. **a**, Response properties of chinchilla SGNs to sinusoidal current injection (50 Hz, amplitude, 0.2 nA) and cell-body sinusoidal mechanical displacement (50 Hz; amplitude, 0.3 μm), and overlap of the two stimuli at ∼90° phase angle. b, Cycle histograms of the same neuron for four relative phase angles, showing single-mode distributions with small changes in the response phase. c, Example of chinchilla SGN exhibiting peak-splitting in the cycle histogram during combined cell-body mechanical displacement and current injection using 50 Hz sinusoidal stimuli. d, Cycle histograms with two modes were obtained at 90° and 180° phase angles.

**Figure S10.**
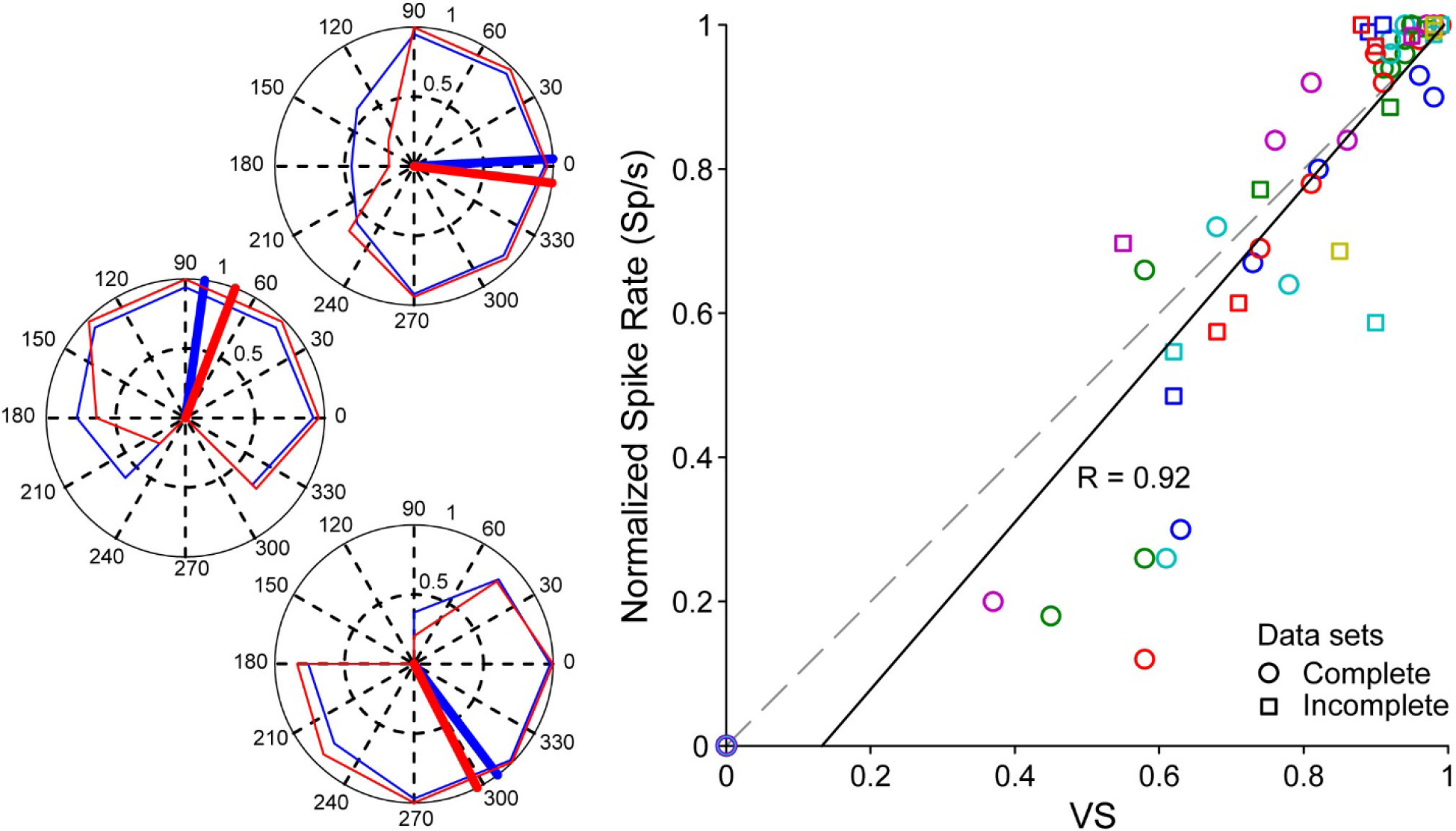
Additional examples of the relationship between firing rate and VS for different relative phase angles between mechanical and current stimulation. Left: polar plots for 3 neurons in which a full circle of relative phase angles was tested. Conventions as in Fig. 3. Right: the relationship between spike rate and VS for all tests, including incomplete datasets. Symbols with circles indicate complete measurement series (n=5) (0°-315° in steps of 45°) with different colors indicating different experiments. Square symbols are from incomplete phase sets (n=6, containing 3 or 4 phase measurements). The spike rates are normalized to the maximum within each series. The dashed line indicates unity, and the black line is the least-square fit, with R the Pearson correlation.

**Figure S11.**
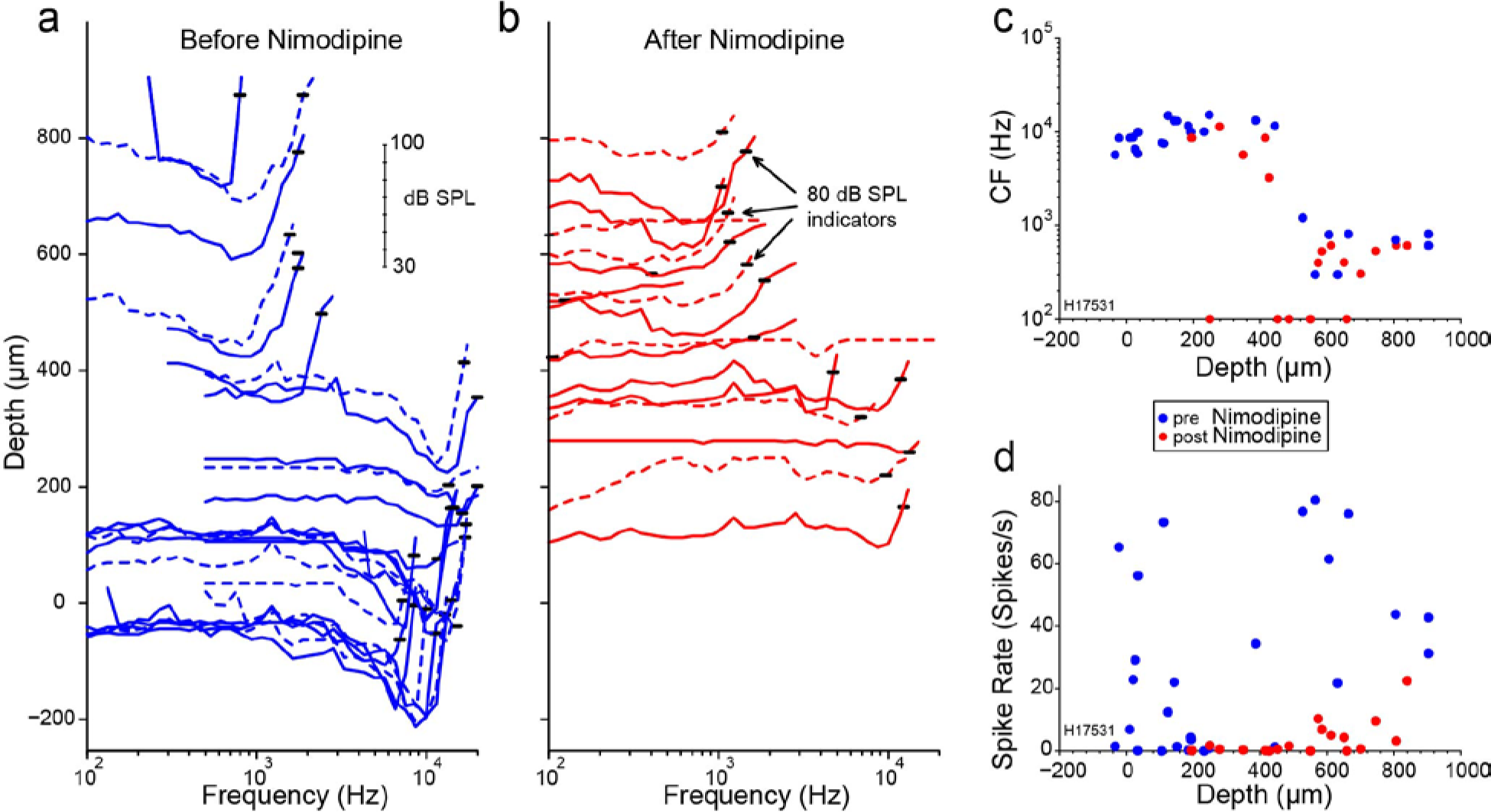
Influence of applied toxins on AN-Fiber responses. Layout as for Fig. 4, but using the Ca^2+^ channel blocker nimodipine. Data from chinchilla.

## Notes

### Competing Interest Statement

The authors have declared no competing interest.

